## Supplementary Information for "Induced fit with replica exchange improves protein complex structure prediction"

Figs. S1 to S16
Tables S1 to S2
SI References

### Supplementary Results

**ReplicaDock 2.0 samples structures from a broad energy-biased population.** To sample aggressively with replica exchange methods, it is necessary to have a significant overlap between energy distributions of replicas. For biomolecular systems, replica exchange utilizes multiple temperature replicas to allow for overlaps, however, it was demonstrated by Zhang *et al.*(1) that small number of temperature replicas can evidently produce broad energy distributions for protein docking sufficient for sampling. With ReplicaDock 2.0 algorithm, we establish three replicas such that the highest temperature replica reflects an unbound state sampling larger conformational changes, whereas the lowest temperature replica reflects a bound state penalizing deleterious backbone conformations. **Fig.S1** compares the energy distribution of RosettaDock 4.0 with ReplicaDock 2.0 for docking target 2CFH, trafficking protein particle complex subunits, 1.55 Å $\mathrm{RMSD}_{BU}$) (2). Thermodynamically, the high-temperature replicas with non-binding conformations have relatively higher energy scores (i.e. poorer binding energy, signifying their unbound nature, as evident in **Fig.S1**, for ReplicaDock 2.0 *in red*). As we go towards low temperature replicas, the energy scores start decreasing (equivalent to better thermodynamic Gibbs free energy, highlighting better binding). The inverse temperatures employed in our study resulted in sampling populations with broader energy distributions with considerable overlap between all three replicas. This enabled the replica exchange method to achieve efficient exchange rates and sample the low-energy structures in the conformational energy landscape.

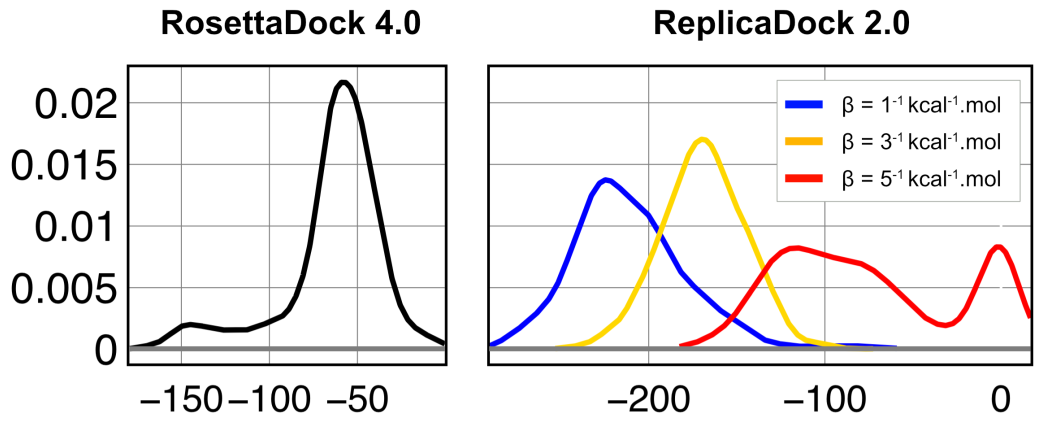


**Fig. S1. Energy distribution** of conformations sampled with RosettaDock 4.0 and ReplicaDock 2.0 (at respective inverse temperatures) for protein target 2CFH

**Global Docking Performance.** To assess global docking performance with ReplicaDock, we performed a rigid-body search with longer MC trials ($10^6 - 10^8$ steps) to identify protein binding interfaces. The protein partners were randomized before each global docking simulation and the protocol generated ∼24,000 candidate structures for each target. With ReplicaDock 2.0, we tested 10 protein targets with only rigid-body translation (2Å) and rotation (4°) starting from a randomly perturbed, initial, docking pose. We observed that for almost all the targets, we produced sub-10 Å structures. We compared ReplicaDock 2.0 global docking performance with ClusPro. ClusPro is a fast-fourier transform (FFT) based method with successful automated server performance in CAPRI. Each ClusPro run identifies low energy structural models which undergo clustering and energy minimization (with CHARMM potential). We obtained these energy-minimized structures from ClusPro, performed Rosetta Relax (energy-minimization) to obtain the clusters. **Fig.S2** demonstrates the comparison of the docking decoys generated with ReplicaDock 2.0

superimposed by the ClusPro clusters (*in red*). Across different levels of flexibility, ReplicaDock 2.0 could produce sub-5 Å models for 8 out of 10 cases. On other hand, ClusPro identified sub-5 Å structures in 6 out of 10 cases, however for most targets, ReplicaDock 2.0 generated lower energy funnels in the near-native region.

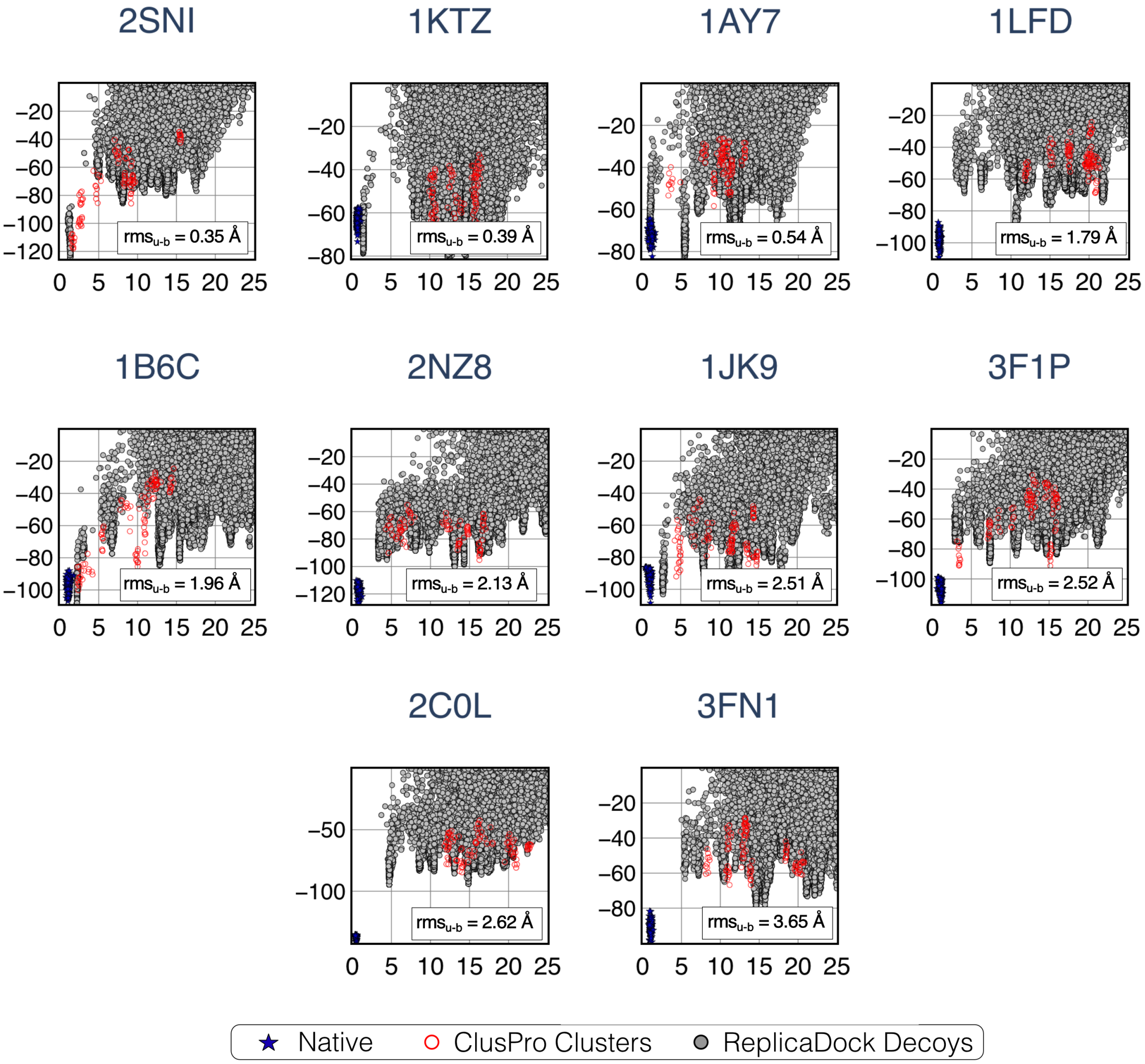


**Fig. S2. Global docking performance.** Interface score (REU) vs I-rmsd (Å) for each of the 10 benchmark targets, arranged by target difficulty. ReplicaDock 2.0 decoys colored in gray and ClusPro models, relaxed with Rosetta and scored with MUDS, highlighted in red.

**Comparison with RoseTTAFold and AlphaFold2.** Development of deep-learning approaches such as AlphaFold (3, 4) and RoseTTAFold (5) have enabled highly accurate three-dimensional structure prediction. Recent methods have extrapolated these sequence-to-structure tools for protein complex prediction, either by incorporating a gap in the protein sequence to indicate a chain-break, or by utilizing paired MSAs. Bryant et al.(6) performed a thorough assessment of AlphaFold and RoseTTAFold on a test set (derived from Dockground Benchmark set 4 (7)) and reported that AlphaFold can predict the structure of hetero-dimeric complexes for up to 59% of the targets. The recent release of AlphaFold-multimer (4), along with thorough assessment (8) of AlphaFold on the Dockground Benchmark set has demonstrated that AlphaFold can produce a DockQ score $\geq 0.23$ for 67% of the hetero-dimeric complexes(4). However, as protein-docking involves prediction of conformational changes from an unbound state of a protein partner to a bound state, elaborate analysis on the predicting structures corresponding to both bound and unbound forms, as well as the possibility of the inclusion of the bound monomers in AlphaFold's training set are unclear. To perform a relatively fair assessment, we tested the performance of AlphaFold (and AlphaFold-multimer) on recent blind CAPRI targets T164 and T165(9). For CAPRI target T164, a structural maintenance of chromosomes flexible hinge domain-containing (SMCHD1) protein complex (PDB ID: 6N64(10)), with an available homologous template (PDB ID: 1GXK, $iRMSD_{template-bound} = 3.84$Å), the predictions by the leading predictors along with the predictions by ReplicaDock2.0, and the deep-learning methods AlphaFold and RoseTTAFold are reported in **Fig.S3,S7**. ReplicaDock2.0 models were initiated from a model created with the homologous template and the best sampled model was reported at an iRMSD of 3.75 Å. We failed to obtain any structures within 5 Å iRMSD with RoseTTAFold, however, with AlphaFold, the best sampled model had an iRMSD of 3.7 Å. For target T164, AlphaFold and AlphaFold-multimer predictions were almost identical (less than 0.5 Å all-atom RMSD difference). Further, we also found that by equipping ReplicaDock2.0 flexible backbone docking with the template from AlphaFold-multimer, we predicted models under 3 Å iRMSD, improving the docking performance by nearly 1 Å (**Fig.S8**). A detailed analysis of AlphaFold models on the bound and unbound templates has been performed and illustrated in **Fig.S7**. For target T165, a monoclonal Antibody bound to varicella-zoster virus glycoprotein (PDB ID: 6vn1(11)), the performance by the deep-learning methods was underwhelming. Although, we could obtain individual monomers with fairly acceptable accuracy (RMSD $<$ 4 Å), predicting the complex structure was challenging and neither of the two methods could identify the appropriate binding orientation or the quaternary structure of the complex. For this case, ReplicaDock2.0 obtained acceptable targets with 4 Å iRMSD (**Fig.S9**).

### Supplementary methods

**Benchmark Sets.** The benchmark sets were collated from Docking Benchmark 5.0(12). First, to test the optimum settings for backbone flexibility, we assembled a smaller, representative, benchmark set of 12 targets (four targets each from rigid, medium and difficult sets). Second, a larger, docking benchmark set comprising all 34 difficult and 44 medium targets from the Docking Benchmark 5.0(12), along with 10 rigid targets. (For target 3R9A, we could not generate RosettaDock4.0 ensembles in reasonable time, and so have excluded it from the benchmark set). Each PDB structure was culled of all HETATM lines, water and non-canonical amino acids. The native-like structures (*in blue*) were produced by performing refinement with small translations (0.1 Å) and rotations (0.5°) on the bound crystal structures. To generate the docking decoys for the CS-based RosettaDock, we ran the protocol described in Marze *et al.*(13) on unbound complexes. For IF-based ReplicaDock, we ran the protocols described further.

**Rosetta Movers used in ReplicaDock 2.0 protocol.** ReplicaDock 2.0 algorithm employs multiple Rosetta movers to perform backbone conformational sampling and docking. The default set-up for ReplicaDock 2.0 involves a replica exchange procedure across three replicas, with inverse temperatures set to $1.5^{-1}$ $kcal^{-1}$.mol, $3^{-1}$ $kcal^{-1}$.mol and $5^{-1}$ $kcal^{-1}$.mol, respectively. 8 trajectories with 3 temperature replicas are run for 2.5 x $10^5$ MC steps and snapshots (structures and temperature/acceptance statistics) are stored after every 1000 steps. The choice of temperature levels for the replicas is dependent on good exchange rates ( 25%). ReplicaDock 2.0 builds on the earlier framework of ReplicaDock protocol of the Rosetta software suite.

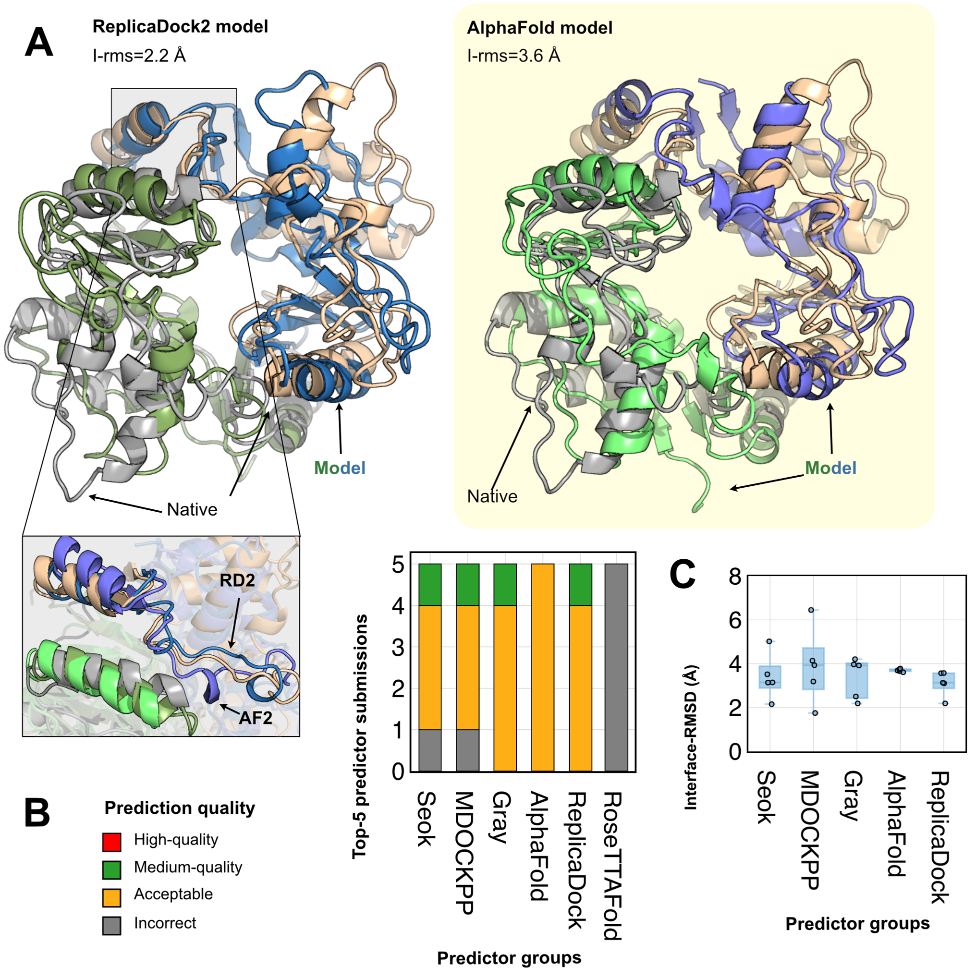


**Fig. S3. ReplicaDock2 and AlphaFold performances on blind CAPRI target T164.** (A) ReplicaDock2.0 prediction (green-blue) superimposed over the bound structure (in tan). Comparison of interface regions between for bound (tan) with ReplicaDock2.0 model (green-blue) and AlphaFold (pale green-pale blue) highlighted below. AlphaFold prediction (pale green-pale blue) superimposed over the bound structure. Binding orientation correlates with the wildtype, but prediction derives mostly from the unbound templates. AlphaFold-multimer was used via available jupyter notebook, but the models generated with AlphaFold-multimer were almost identical (less than 0.5 Å all-atom RMSD difference) to the models produced with AlphaFold. (B) Top-5 predictions for the predictor groups colored by the prediction quality. AlphaFold and ReplicaDock2 capture better quality decoys than predictions. RoseTTAFold predictions are inaccurate. (C) Interface-RMSD (Å) for the top-5 predictions by the predictor groups. RoseTTAFold models could not capture models within 8 Å and are excluded from this plot.

The module is currently available via `RosettaScripts` interface and can be combined with alternative movers developed as part of the `ThermodynamicMover` class. The description of each mover employed in the ReplicaDock 2.0 algorithm is discussed further:

**EncounterConstraintMover.** With increasing temperatures in the higher replicas, there is a possibility that the protein partners can diffuse away from each other. To prevent this non-bound diffusion of the protein partners, we employ an encounter constraint. Encounter constraint is implemented as a flat-bottom distance restraint to restrain the protein partners within a certain range of each other. The constraint penalizes conformations if they are further than $l_{max} = \mathrm{g} + \mathrm{d}_1 + d_2$ apart, such that $g$ represents the input gap parameter (set to 8 Å) and $d_i$ denotes the distance between the farthest surface $C_\alpha$ atoms of partner $i$ to its center of mass. The penalty is enforced with a harmonic function ($\mathrm{H}(l) = \mathrm{k}(l - l_{\mathrm{max}})$) for all $l > l_{max}$.

**DockingInitialPertubationMover.** For each trajectory, the protein partners are first brought into contact with an initial perturbation. This perturbation is produced by the `DockingInitialPerturbation` mover. For global docking, we can initialize our docking trajectories by randomizing the protein partners around each other (set `randomize1` and `randomize2` options to true). This allows the protocol to completely lose the context of any local binding interface and perform blind, global docking. For local docking, we preserve local binding site information, and instead slide the protein partners into contact to produce our initial docking pose (set `slide` to true). Sliding the initial docking pose into contact also allows the protocol to identify putative binding interface and select residues to sample backbone degrees of freedom.

**RigidBodyPerturbNoCenter.** To dock the protein partners in the rigid-body space i.e. with respect to each other, we use this mover to perform random rotations and translations to the protein partners in the docking pose. For local docking with ReplicaDock 2.0 algorithm, the magnitudes of the random translation and rotation are set to 1 Å and 1° respectively. We expand these to translational magnitudes of 2 Å and rotational magnitudes of 2° for global docking runs.

**HamiltonianExchangeMover.** The HamiltonianExchangeMover is an update to the ParallelTempering-Mover in Rosetta and allows modulating the temperatures as well as the score functions within the replicas. This mover is governed by a configuration file (input by the option `temp_file` and provides the mover with the temperature levels as well as the score-functions across each temperature level. The temperature information is fed via inverse temperatures and score-function terms can be modulated independently. Here, we have set the inverse temperatures as described earlier and scale down the repulsive energy potential weights for the higher replicas (i.e. for medium-temperature and hot-temperature replicas with inverse temperatures $3^{-1}$ kcal$^{-1}$.mol and $5^{-1}$ kcal$^{-1}$.mol, we bring down the repulsive weight to 0.75 and 0.5 respectively). With the reduction in repulsive weights, our aim is to allow generation of those backbone conformations which would be otherwise rejected, thereby enabling aggressive sampling.

**BackrubMover.** For enabling backbone sampling, we employ the backrub mover over the selected interface residues. Two terminal residues for each contiguous fragment are selected as pivots and local moves using backbone dihedral rotations around their axes is performed with backrub. The maximum angular displacement for a fragment between the pivot residues can be manipulated by the user to make sampling more or less aggressive as necessary. The mobile residues are selected via a Rosetta `movemap` defined with `RESIDUESELECTORS` in Rosetta, and input via the `MoveMapFactory`. This allows the dynamic allocation of the residues for each docking pose and results in an on-the-fly backbone sampling dependent on sampled protein interface in a pose. For directed induced-fit, we override this behaviour by providing the `BackrubMover` with the pivot residues as a list (this is input by the `pivot_residues` option).

**MetropolisHastingsMover.** The `MetropolisHastingsMover` serves as the container that performs all the mover operations listed above with MC sampling. For the given temperatures of the replicas, the backbone and docking movers, this mover performs the trial MC moves and also records statistics of acceptance post each trial move. We pass the `HamiltonianExchangeMover`, `RigidBodyPerturbNoCenter` and `BackrubMover` as submovers within the `MetropolisHastingsMover` and adjust the sampling frequency of each mover by modulating the `sampling_weight` option. With the `HamiltonianExchangeMover` as the temperature controller and the `MetropolisHastingsMover` as the sampler, we perform replica exchange with backbone moves and docking in ReplicaDock 2.0.

**Motif Updated Dock Score (MUDS) optimization.** Prior work by Marze *et al.*(13) introduced the Motif Dock Score (MDS) for low-resolution protein docking. This score term was created by first culling protein structures with one or more interfaces from the Protein Data Bank (PDB) (14, 15) and obtaining their all-atom Rosetta energies. Then a discrete score-grid is initialized with the energies of the interacting residue pairs, if the interacting pair score is below an energy threshold (generally 0 REU) and the interacting residues have their C$\beta$ atoms within 10 Å of each other. MDS was built only on inter-chain residue pairs and was developed to be a fast, low-resolution equivalent of all-atom interface energies. A caveat of MDS, as reported by Marze *et al.*(13) is the inability of MDS to penalize clashing or high-energy residue-residue interactions. This was overcome by incorporating a Van der Waals clash penalty term. Since in ReplicaDock 2.0, the protocol aims to sample backbone degrees of freedom, by not incorporating single-body terms, the backbone sampling was often misdirected resulting in substantial intra-chain steric clashes and unrealistic protein backbones. To circumvent around this issue, we incorporated backbone statistic terms which are majorly dependent on the backbone heavy atoms of a protein pose. These terms incorporated the likelihood of observing the rotamers in a certain $\phi$ or $\psi$ angles (`p_aa_pp`), Ramachandran score to determine whether a residue fits in the Ramachandran space (`rama_prepro`) and a penalty towards deviation from the $\omega$ dihedrals(`omega`). We optimized MUDS with different weights and found that weighing the individual score terms as determined for the all-atom score function resulted in more acceptable CAPRI-quality decoys in the top-10% scoring structures (**Fig.S6**). The exact formulation employed for MUDS is as follows:

`muds_2021 = 1.0*motif_dock + 0.55*fa_rep + 1.0*fa_atr + 0.5*rama_prepro + 0.48*omega + 0.61*_aa_pp`

**Docking Metrics.** To evaluate the docking performance for the docking metrics, we used standard CAPRI metrics. These metrics are defined here as follows:

**Interface Score** : The Interface score (*in Rosetta Energy Units, REU*) is a Rosetta equivalent of the thermodynamic binding energy upon protein association. This score is estimated by calculating the total score of the protein receptor-ligand complex and then by subtracting the individual (monomeric) scores of protein receptor and ligand pulled apart.

$$\Delta\Delta G_{\text{interface}} = \Delta G_{\text{AB}} - \Delta G_{\text{A}} - \Delta G_{\text{B}}$$

**C$_\alpha$ RMSD** : The C$_\alpha$ root mean square deviation (RMSD) of the candidate structure with the experimental crystal structure.

**Interface-RMSD(Irms)** : The RMSD over all the backbone atoms of the residues on the interface in a docked protein structure to the experimental bound crystal structure. Interface residues are defined as those residues that lie within 10 Å of any residue on the other binding partner.
**RMSD** : The heavy atom RMSD between the atoms of the candidate structure and the experimental crystal structure. The backbone RMSDs of the monomers also employ this metric.
**⟨N5⟩** : The number of near-native decoys in the 5 top-scoring structures reported by Interface score. This value ranges from 0 to 5, with 0 (worst) being no near-native models were generated in the 5 top-scoring structures, and 5 (best) implying all top-scoring structures were near-native. A model is attributed as a near-native model if its CAPRI quality is acceptable or better. We average the N5 metric by bootstrapping 1,000 random samples for robustness.

**Scripts and tutorials.** The unbound monomers (protein receptor and ligand structures) are oriented roughly 10-15 Å apart each other, briefly defining the binding interface. This docking poses is passed on for the first stage of ReplicaDock2.0 simulations, that involves replica-exchange and backbone sampling. To run ReplicaDock2.0 protocol, message passing interface (MPI) mode is essential. The number of processors that have to be used are calculated with the following formula: `num(processors) = nstruct`×`n_replica + 2` , where `n_replica` is the number of temperature replicas, 2 denotes the processors for the job distributor and file I/O, and `nstruct` denotes any positive integer.

Rosetta executables must be compiled with `extras=mpi` for running ReplicaDock 2.0 simulations. The trajectory is output `trials.stats` file that records the acceptance rates of each temperature level. Generated decoys are tagged with the temperature level, the trajectory number and the snapshot number within the trajectory. The following XML script is the script employed for the benchmarking of ReplicaDock 2.0 protocol. The backrub motions can be tuned to perform more or less aggressive motions as necessary by the user. For rigid-body global docking, the `Backrub` mover can be commented out, and the `DockingInitialPerturbation` mover could be modified to randomize the inputs. For directed induced-fit, the Backrub mover is modified to include pivot-residues. The `sampling_weight` option in the Metropolis-Hastings mover container allows to tune the operation of any mover within it. Here, we provide an example of the XML scripts and the flags used to benchmark the protocol. More demos/tutorials highlighting all the interface residue test selections, global docking scripts and directed-induced fit are available in the https://www.rosettacommons.org/, in `demos/public/replicadock2` directory. The protocol runs with the RosettaScripts executable:

```
rosetta_scripts.mpi.linuxgccrelease -np X @flags_replica_dock
```

The description of the necessary files is defined further.

`flags_replica_dock`

| Flag | Description | Input |
|---|---|---|
| **in:file:s** | Name of the input PDB file (initial structure of the docking complex) | P.pdb |
| **in:file:native** | Name of the native PDB file | native.pdb |
| **partners** | To define docking partners with their chain IDS, for e.g: A_B, A_HL, AB_CD | A_B* |
| **parser:protocol** | The XML script that has details of the protocol to be run | replicadock.xml |
| **evaluation:DockMetrics** | To compute all the docking metrics (Irms, L-rms, $f_{nat}$ ,etc) for replica exchange docking | true |
| **score:weights** | The score-function and its respective weights from the weights file | muds_2021 |
| **run:n_replica** | The number of replicas to run in replica docking | 3 |
| **out:nstruct** | The number of trajectories to run with replica docking | 8 |
| **out:path:all** | The output path for all generated files | output |
| **out:file:silent** | The output path for all the silent (.out) files | decoys.out |
| **out:file:scorefile** | The output score file to be generated with the protocol | scores.sc |
| **out:mpi_tracer_to_file** | The output file that contains all the tracer output | tracer.out |
| **multiple_processes_writing _to_one_directory** | Flag option to indicate that MPI mode is in progress and multiple processors would contribute to the same score file | True |

`hamiltonians_cen.txt`

```
GRID_DIM 1
GLOBAL_PATCH atom_pair_constraint = 5
1 1.5 muds_2021 NOPATCH fa_rep *= 1.0
2 3.0 muds_2021 NOPATCH fa_rep *= 0.75
3 5.0 muds_2021 NOPATCH fa_rep *= 0.5
#ETABLE FA_STANDARD_SOFT10 fa_rep *= 1.1
```

`replicadock.xml`

```
<ROSETTASCRIPTS>
<SCOREFXNS>
    <ScoreFunction name="score_dock_low" weights="muds_2021"/>
    <ScoreFunction name="score_analyze" weights="motif_dock_score"/>
</SCOREFXNS>
<FILTERS>
</FILTERS>
<RESIDUE_SELECTORS>
    <Chain name="chA" chains="A" />
    <Chain name="chB" chains="B" />
    <Neighborhood name="chA_neighbours" selector="chA" distance="8.0"/>
    <Neighborhood name="chB_neighbours" selector="chB" distance="8.0"/>
```

```
    <And name="interfaceA" selectors="chB_neighbours,chA"/>
    <And name="interfaceB" selectors="chA_neighbours,chB"/>
    <Or name="interfaceAB" selectors="interfaceA,interfaceB"/>
</RESIDUE_SELECTORS>
<MOVE_MAP_FACTORIES>
    <MoveMapFactory name="Interface">
            <Backbone residue_selector="interfaceAB"/>
            <Chi residue_selector="interfaceAB"/>
    </MoveMapFactory>
</MOVE_MAP_FACTORIES>
<SIMPLE_METRICS>
</SIMPLE_METRICS>
<MOVERS>
    <!-- setup jumps and constraints-->
    <DockSetupMover name="setup_jump"/>
    <AddEncounterConstraintMover name="encounter_cst" gap="8" />
    <RigidBodyPerturbNoCenter name="rb_mover" rot_mag="1" trans_mag="1"/>
    <HamiltonianExchange name="h_exchange" temp_file="hamiltonians_cen.txt" temp_stride="1000" stats_file="tempering.stats"/>
    <DockingInitialPerturbation name="init_pert" slide="1" />
    <!--DockingInitialPerturbation name="init_pert" randomize1="1" randomize2="1" /--> <!-- Uncomment this for global docking-->
    <TrialCounterObserver name="count" file="trial.stats"/>
    <SilentTrajectoryRecorder name="traj" score_stride="1" stride="1000" cumulate_replicas="1" />
    <Backrub name="bbmover" movemap_factory="Interface" require_mm_bend="0"/>
    <!--Backrub name="bbmover" pivot_residues="" max_angle_disp_4="3.1" max_angle_disp_7="3.0" max_angle_disp_slope="-0.5" require_mm_bend="0"/--> <!-- Additional tags to perform aggressive BB sampling or to input specific pivot residues while docking-->
    <MetropolisHastings name="sampler" trials="250000" scorefxn="score_dock_low" > <!--trial number normally use 10^6-10^8 for productive global docking simulation-->
        <Add mover_name="h_exchange"/>
        <Add mover_name="traj"/>
        <Add mover_name="count"/>
        <Add mover_name="rb_mover"/>
        <Add mover_name="bbmover" sampling_weight="2"/>
    </MetropolisHastings>
 </MOVERS>
 <PROTOCOLS>
     <Add mover_name="setup_jump"/>
     <Add mover_name="encounter_cst"/>
     <Add mover_name="init_pert"/>
     <Add mover_name="sampler"/>
</PROTOCOLS>
</ROSETTASCRIPTS>
```

### Path to motif_dock_score tables

```
-mh:path:scores_BB_BB Rosetta/main/database/additional_protocol_data/motif_dock/xh_16_
-mh:score:use_ss1 false
-mh:score:use_ss2 false
-mh:score:use_aa1 true
-mh:score:use_aa2 true
```

### Supplementary Tables

**Table S1. Comparison of leading docking methods with ReplicaDock 2.0 (derived from Marze *et al.*(13)).** (1) Nearest-native structures from rigid-body docking selected for refinement. (2) Half successes awarded for targets with multiple binding sites evaluated, where at least one but not all binding sites are captured. (3) 2.5 Å cutoff for near-native structures. (4) Cases where bootstrapping gives ≥50% chance of N5≥3 are considered successfully docked. (5) For CAPRI sets, medium and difficult targets are combined, comprising all targets without at least one high-quality prediction by any predictor. (6) Lensink *et al.*(16) (7) Hwang *et al.*(17) (8) Vreven *et al.*(12)

| **Methods** | | | | | | **Performance** | | |
|---|---|---|---|---|---|---|---|---|
| Method | Description | Flexibility? | Benchmark Set | Docking Search | Success Metric | Easy Targets | Medium Targets[5] | Difficult Targets[5] |
| HADDOCK (2017) | Restraint-based docking, minimization | Yes | CASP-CAPRI[6] | Mixed global/local | N10 = 1 | 12/12 (100%) | 4/13 (31%) | |
| ClusPro (2017) | FFT docking, cluster evaluation | No | CAPRI Rds. 13−35 | Mixed global/local | N10 = 1 | 12.5[2]/16 (78%) | 6.5[2]/26 (25%) | |
| iATTRACT (2015) | Rigid-body docking, interface refinement | Yes | Docking Benchmark 4.0[7] | Global[1] | N200 = 30 | 55/119 (46%) | 9/28 (32%) | 0/19 (0%) |
| ZDOCK (2011) | FFT docking, model evaluation | No | Docking Benchmark 4.0[7] | Global | N100 = 1[3] | 58/121 (48%) | 7/30 (23%) | 0/25 (0%) |
| Rosetta Dock 3.2 (2011) | Monte Carlo docking, model evaluation | Yes | Docking Benchmark 4.0[7] | Local | N5 = 3 | 49/84 (58%) | 5/17 (29%) | 2/14 (14%) |
| RosettaDock 4.0 (2018) | Monte Carlo docking, model evaluation | Yes | Docking Benchmark 5.0[8] | Local | N5 = 3[4] | 10/13 (77%) | 21/43 (49%) | 10/32 (31%) |
| ReplicaDock 2.0 (2021) | Replica Exchange Monte Carlo docking | Yes | Docking Benchmark 5.0[8] | Local | N5 > 3[4] | 8/10 ( 80%) | 27/44 (61%) | 12/34 (35%) |

**Table S2. Performance of RosettaDock 4.0 vs. ReplicaDock 2.0 across an 88-target benchmark set. 5,000 decoys were generated by each protocol for each target. Bootstrapped N5 values (plus standard deviations), both after the low-resolution phase and after the full protocol, are listed for each target. Success is defined as $\langle N5 \rangle \geq 3$ for the N5 metrics.**

| Target | Difficulty | Average N5 | | | |
|---|---|---|---|---|---|
| | | RosettaDock (LowRes) | RosettaDock | ReplicaDock (LowRes) | ReplicaDock |
| 1AY7 | Rigid | 0.6 ± 0.8 | 5.0 ± 0.0 | 5.0 ± 0.0 | 5.0 ± 0.0 |
| 1BVK | Rigid | 0.0 ± 0.0 | 4.9 ± 0.5 | 3.2 ± 1.1 | 3.6 ± 1.1 |
| 1KTZ | Rigid | 4.3 ± 0.9 | 4.7 ± 0.6 | 5.0 ± 0.0 | 5.0 ± 0.0 |
| 1MAH | Rigid | 1.5 ± 1.2 | 4.7 ± 0.6 | 5.0 ± 0.2 | 5.0 ± 0.0 |
| 1MLC | Rigid | 0.0 ± 0.2 | 0.1 ± 0.3 | 0.0 ± 0.0 | 0.0 ± 0.0 |
| 2BTF | Rigid | 5.0 ± 0.3 | 5.0 ± 0.0 | 5.0 ± 0.0 | 0.6 ± 0.9 |
| 2JEL | Rigid | 3.7 ± 1.3 | 4.5 ± 0.8 | 5.0 ± 0.0 | 5.0 ± 0.0 |
| 2PCC | Rigid | 0.0 ± 0.0 | 3.0 ± 1.4 | 5.0 ± 0.0 | 5.0 ± 0.0 |
| 2SIC | Rigid | 0.6 ± 0.8 | 5.0 ± 0.0 | 5.0 ± 0.0 | 5.0 ± 0.0 |
| 2SNI | Rigid | 4.1 ± 0.9 | 5.0 ± 0.0 | 5.0 ± 0.0 | 5.0 ± 0.0 |
| 1B6C | Medium | 5.0 ± 0.0 | 5.0 ± 0.0 | 5.0 ± 0.1 | 5.0 ± 0.0 |
| 1CGI | Medium | 1.9 ± 1.3 | 1.9 ± 1.4 | 5.0 ± 0.0 | 3.0 ± 1.1 |
| 1FC2 | Medium | 0.0 ± 0.2 | 0.0 ± 0.2 | 0.2 ± 0.5 | 3.1 ± 1.3 |
| 1GP2 | Medium | 1.6 ± 1.1 | 1.6 ± 1.1 | 0.0 ± 0.0 | 0.0 ± 0.0 |
| 1GRN | Medium | 1.4 ± 1.1 | 1.3 ± 1.0 | 5.0 ± 0.0 | 5.0 ± 0.0 |
| 1HE8 | Medium | 4.5 ± 0.7 | 4.6 ± 0.7 | 5.0 ± 0.0 | 5.0 ± 0.0 |
| 1I2M | Medium | 0.0 ± 0.0 | 0.0 ± 0.0 | 5.0 ± 0.0 | 4.6 ± 0.8 |
| 1IB1 | Medium | 0.0 ± 0.0 | 0.0 ± 0.0 | 0.0 ± 0.0 | 0.0 ± 0.0 |
| 1IJK | Medium | 3.7 ± 1.1 | 3.7 ± 1.0 | 5.0 ± 0.0 | 5.0 ± 0.0 |
| 1JIW | Medium | 0.8 ± 1.1 | 0.9 ± 1.1 | 0.1 ± 0.3 | 2.5 ± 1.2 |
| 1K5D | Medium | 0.9 ± 0.9 | 0.9 ± 0.9 | 0.0 ± 0.1 | 1.2 ± 1.0 |
| 1KKL | Medium | 3.1 ± 1.2 | 3.2 ± 1.2 | 5.0 ± 0.0 | 0.0 ± 0.0 |
| 1LFD | Medium | 5.0 ± 0.0 | 5.0 ± 0.0 | 5.0 ± 0.0 | 5.0 ± 0.0 |
| 1M10 | Medium | 0.0 ± 0.2 | 0.0 ± 0.2 | 0.2 ± 0.5 | 3.0 ± 1.2 |
| 1MQ8 | Medium | 5.0 ± 0.0 | 5.0 ± 0.0 | 0.0 ± 0.0 | 2.3 ± 1.3 |

| | | | | | | | | | | | | | |
|---|---|---|---|---|---|---|---|---|---|---|---|---|---|
| 1N2C | Medium | 1.3 | ± | 0.8 | 0.0 | ± | 0.0 | 5.0 | ± | 0.0 | 0.1 | ± | 0.3 |
| 1NW9 | Medium | 3.6 | ± | 1.1 | 3.6 | ± | 1.1 | 0.0 | ± | 0.0 | 1.4 | ± | 1.1 |
| 1R6Q | Medium | 2.7 | ± | 1.3 | 2.7 | ± | 1.3 | 5.0 | ± | 0.0 | 5.0 | ± | 0.0 |
| 1SYX | Medium | 5.0 | ± | 0.0 | 5.0 | ± | 0.0 | 5.0 | ± | 0.0 | 5.0 | ± | 0.0 |
| 1WQ1 | Medium | 2.8 | ± | 1.3 | 2.8 | ± | 1.3 | 5.0 | ± | 0.0 | 4.6 | ± | 0.6 |
| 1XQS | Medium | 4.2 | ± | 0.9 | 4.2 | ± | 0.8 | 5.0 | ± | 0.0 | 4.3 | ± | 0.8 |
| 1ZM4 | Medium | 2.5 | ± | 1.1 | 2.5 | ± | 1.2 | 0.5 | ± | 0.9 | 4.9 | ± | 0.3 |
| 2CFH | Medium | 5.0 | ± | 0.0 | 5.0 | ± | 0.0 | 5.0 | ± | 0.0 | 5.0 | ± | 0.0 |
| 2H7V | Medium | 2.6 | ± | 1.1 | 2.6 | ± | 1.2 | 5.0 | ± | 0.0 | 3.0 | ± | 1.1 |
| 2HRK | Medium | 4.4 | ± | 0.8 | 4.4 | ± | 0.9 | 1.2 | ± | 1.0 | 4.9 | ± | 0.4 |
| 2NZ8 | Medium | 3.6 | ± | 1.1 | 3.7 | ± | 1.1 | 3.2 | ± | 1.1 | 3.6 | ± | 1.1 |
| 2OZA | Medium | 0.0 | ± | 0.0 | 0.0 | ± | 0.0 | 0.0 | ± | 0.0 | 0.0 | ± | 0.0 |
| 2Z0E | Medium | 0.9 | ± | 0.9 | 0.9 | ± | 0.9 | 0.1 | ± | 0.5 | 0.1 | ± | 0.4 |
| 3AAA | Medium | 1.5 | ± | 1.1 | 1.5 | ± | 1.1 | 0.0 | ± | 0.2 | 0.0 | ± | 0.0 |
| 3AAD | Medium | 0.9 | ± | 0.9 | 0.9 | ± | 0.9 | 0.0 | ± | 0.0 | 0.0 | ± | 0.0 |
| 3BX7 | Medium | 2.4 | ± | 1.4 | 2.5 | ± | 1.3 | 5.0 | ± | 0.0 | 2.6 | ± | 1.2 |
| 3CPH | Medium | 0.4 | ± | 0.7 | 0.4 | ± | 0.7 | 0.7 | ± | 0.9 | 2.1 | ± | 1.2 |
| 3DAW | Medium | 4.4 | ± | 0.8 | 4.4 | ± | 0.8 | 5.0 | ± | 0.0 | 3.1 | ± | 1.1 |
| 3EO1 | Medium | 5.0 | ± | 0.2 | 5.0 | ± | 0.2 | 5.0 | ± | 0.0 | 5.0 | ± | 0.0 |
| 3G6D | Medium | 1.1 | ± | 1.0 | 1.1 | ± | 1.1 | 0.0 | ± | 0.1 | 5.0 | ± | 0.0 |
| 3HI6 | Medium | 0.0 | ± | 0.1 | 0.0 | ± | 0.2 | 0.0 | ± | 0.0 | 0.4 | ± | 0.7 |
| 3L5W | Medium | 5.0 | ± | 0.0 | 5.0 | ± | 0.0 | 5.0 | ± | 0.0 | 3.8 | ± | 1.0 |
| 3S9D | Medium | 0.8 | ± | 0.9 | 3.9 | ± | 1.1 | 0.1 | ± | 0.0 | 4.5 | ± | 0.8 |
| 3SZK | Medium | 3.4 | ± | 1.2 | 3.3 | ± | 1.2 | 5.0 | ± | 0.0 | 5.0 | ± | 0.0 |
| 3V6Z | Medium | 0.0 | ± | 0.0 | 0.0 | ± | 0.0 | 5.0 | ± | 0.0 | 2.1 | ± | 1.2 |
| 4FZA | Medium | 1.0 | ± | 1.0 | 1.0 | ± | 1.0 | 1.6 | ± | 1.2 | 5.0 | ± | 0.0 |
| 4IZ7 | Medium | 0.0 | ± | 0.1 | 0.0 | ± | 0.1 | 0.0 | ± | 0.0 | 0.0 | ± | 0.0 |
| 4JCV | Medium | 2.5 | ± | 1.1 | 2.6 | ± | 1.2 | 2.6 | ± | 1.3 | 5.0 | ± | 0.0 |
| 4LW4 | Medium | 1.6 | ± | 1.1 | 1.5 | ± | 1.1 | 4.7 | ± | 0.8 | 3.7 | ± | 1.3 |

| | | | | | | | | | | | | | |
|---|---|---|---|---|---|---|---|---|---|---|---|---|---|
| 1ACB | Difficult | 2.3 | ± | 1.2 | 2.3 | ± | 1.2 | 5.0 | ± | 0.1 | 3.2 | ± | 1.2 |
| 1ATN | Difficult | 1.9 | ± | 1.1 | 1.9 | ± | 1.1 | 4.7 | ± | 0.6 | 2.5 | ± | 1.2 |
| 1BGX | Difficult | 0.0 | ± | 0.0 | 0.0 | ± | 0.0 | 0.0 | ± | 0.0 | 0.0 | ± | 0.0 |
| 1BKD | Difficult | 0.0 | ± | 0.0 | 0.0 | ± | 0.0 | 0.0 | ± | 0.0 | 0.0 | ± | 0.0 |
| 1DE4 | Difficult | 2.3 | ± | 1.2 | 2.3 | ± | 1.2 | 0.0 | ± | 0.0 | 4.8 | ± | 0.5 |
| 1E4K | Difficult | 0.0 | ± | 0.2 | 0.0 | ± | 0.2 | 0.0 | ± | 0.0 | 0.0 | ± | 0.0 |
| 1EER | Difficult | 0.1 | ± | 0.4 | 0.1 | ± | 0.4 | 5.0 | ± | 0.2 | 0.0 | ± | 0.0 |
| 1F6M | Difficult | 0.3 | ± | 0.7 | 0.2 | ± | 0.6 | 0.0 | ± | 0.0 | 0.0 | ± | 0.0 |
| 1FAK | Difficult | 0.0 | ± | 0.2 | 0.0 | ± | 0.2 | 0.0 | ± | 0.0 | 0.0 | ± | 0.0 |
| 1FQ1 | Difficult | 5.0 | ± | 0.1 | 5.0 | ± | 0.1 | 4.9 | ± | 0.4 | 5.0 | ± | 0.0 |
| 1H1V | Difficult | 0.0 | ± | 0.0 | 0.0 | ± | 0.0 | 0.0 | ± | 0.0 | 0.0 | ± | 0.0 |
| 1IBR | Difficult | 0.0 | ± | 0.0 | 0.0 | ± | 0.0 | 0.0 | ± | 0.0 | 0.0 | ± | 0.0 |
| 1IRA | Difficult | 0.0 | ± | 0.0 | 0.0 | ± | 0.0 | 0.0 | ± | 0.0 | 0.0 | ± | 0.0 |
| 1JK9 | Difficult | 5.0 | ± | 0.0 | 5.0 | ± | 0.0 | 5.0 | ± | 0.0 | 5.0 | ± | 0.0 |
| 1JMO | Difficult | 1.4 | ± | 1.0 | 1.4 | ± | 1.0 | 4.6 | ± | 0.8 | 1.6 | ± | 1.2 |
| 1JZD | Difficult | 1.9 | ± | 1.3 | 1.9 | ± | 1.2 | 5.0 | ± | 0.0 | 5.0 | ± | 0.0 |
| 1PXV | Difficult | 2.6 | ± | 1.3 | 2.5 | ± | 1.3 | 1.3 | ± | 1.1 | 3.2 | ± | 0.6 |
| 1R8S | Difficult | 0.0 | ± | 0.0 | 0.0 | ± | 0.0 | 0.1 | ± | 0.3 | 0.0 | ± | 0.1 |
| 1RKE | Difficult | 0.0 | ± | 0.1 | 0.0 | ± | 0.1 | 0.0 | ± | 0.0 | 1.6 | ± | 1.2 |
| 1Y64 | Difficult | 0.0 | ± | 0.0 | 0.0 | ± | 0.0 | 0.0 | ± | 0.0 | 0.0 | ± | 0.0 |
| 1ZLI | Difficult | 0.8 | ± | 0.9 | 0.7 | ± | 0.8 | 1.2 | ± | 1.0 | 0.0 | ± | 0.0 |
| 2C0L | Difficult | 4.0 | ± | 1.0 | 4.0 | ± | 1.0 | 0.0 | ± | 0.0 | 0.0 | ± | 0.0 |
| 2HMI | Difficult | 2.9 | ± | 1.3 | 2.9 | ± | 1.3 | 0.0 | ± | 0.0 | 0.0 | ± | 0.1 |
| 2I9B | Difficult | 1.0 | ± | 1.0 | 1.1 | ± | 1.0 | 0.1 | ± | 0.4 | 2.2 | ± | 1.2 |
| 2IDO | Difficult | 3.7 | ± | 1.1 | 3.7 | ± | 1.0 | 5.0 | ± | 0.0 | 4.0 | ± | 0.9 |
| 2J7P | Difficult | 0.0 | ± | 0.0 | 0.0 | ± | 0.0 | 0.0 | ± | 0.0 | 0.0 | ± | 0.0 |
| 2O3B | Difficult | 1.4 | ± | 1.1 | 1.3 | ± | 1.1 | 1.4 | ± | 1.4 | 4.0 | ± | 1.0 |
| 2OT3 | Difficult | 0.0 | ± | 0.0 | 0.0 | ± | 0.0 | 5.0 | ± | 0.0 | 0.0 | ± | 0.0 |
| 3AAD | Difficult | 0.0 | ± | 0.0 | 0.0 | ± | 0.0 | 0.0 | ± | 0.0 | 0.0 | ± | 0.0 |

| | | | | | | | | | | | | | |
|---|---|---|---|---|---|---|---|---|---|---|---|---|---|
| 3F1P | Difficult | 5.0 | ± | 0.1 | 5.0 | ± | 0.1 | 5.0 | ± | 0.0 | 5.0 | ± | 0.0 |
| 3FN1 | Difficult | 1.5 | ± | 1.2 | 1.5 | ± | 1.1 | 5.0 | ± | 0.0 | 4.9 | ± | 0.5 |
| 3H11 | Difficult | 5.0 | ± | 0.0 | 5.0 | ± | 0.0 | 5.0 | ± | 0.0 | 5.0 | ± | 0.0 |
| 3L89 | Difficult | 4.3 | ± | 0.8 | 4.4 | ± | 0.8 | 5.0 | ± | 0.0 | 5.0 | ± | 0.0 |
| 4GAM | Difficult | 2.3 | ± | 1.2 | 2.4 | ± | 1.2 | 0.0 | ± | 0.0 | 0.0 | ± | 0.0 |

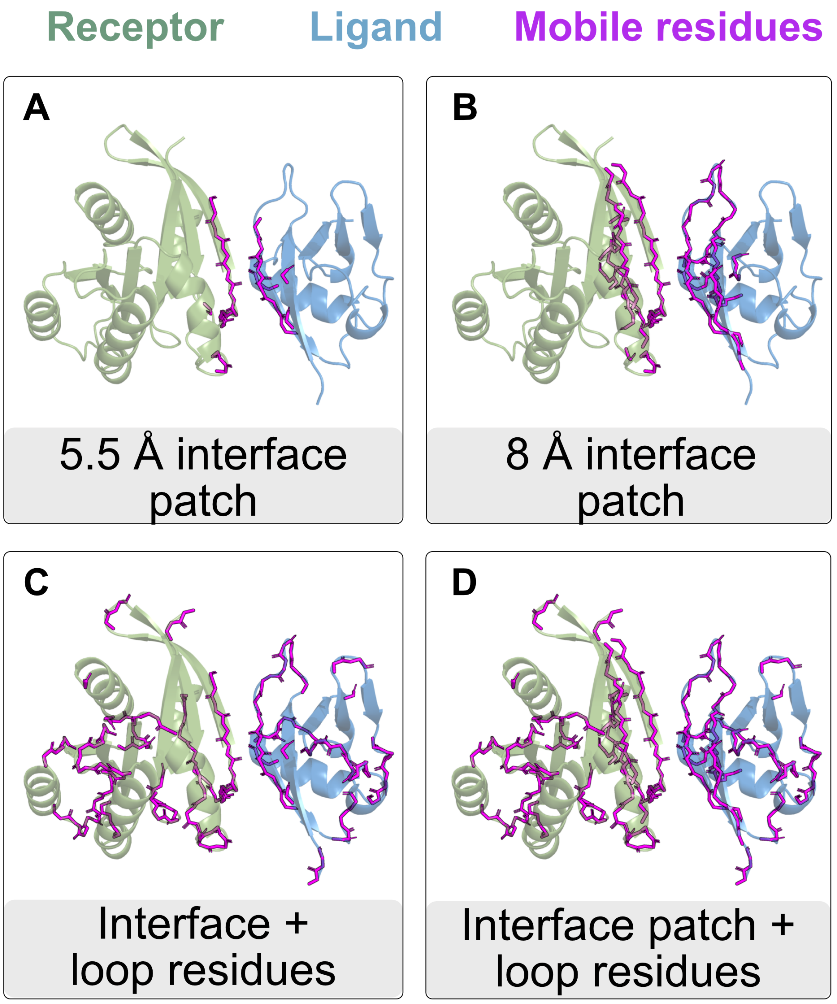


**Fig. S4. Interface residue selections** (*in magenta*) highlighted over a protein target (Receptor, *in green* and ligand, *in blue*). The four residues selections are as follows: (1) 5.5 Å interface patch, (2) 8 Å interface patch (3) 5.5 Å interface patch + loops, (4) 8 Å interface patch + loops. Note that, we also performed a test set by including all the residues of the protein for backbone sampling, however, with T-REMC, such simulations resulted in distortion of the protein quaternary structure (i.e. resulted in protein unfolding). Therefore, we chose to exclude that test.

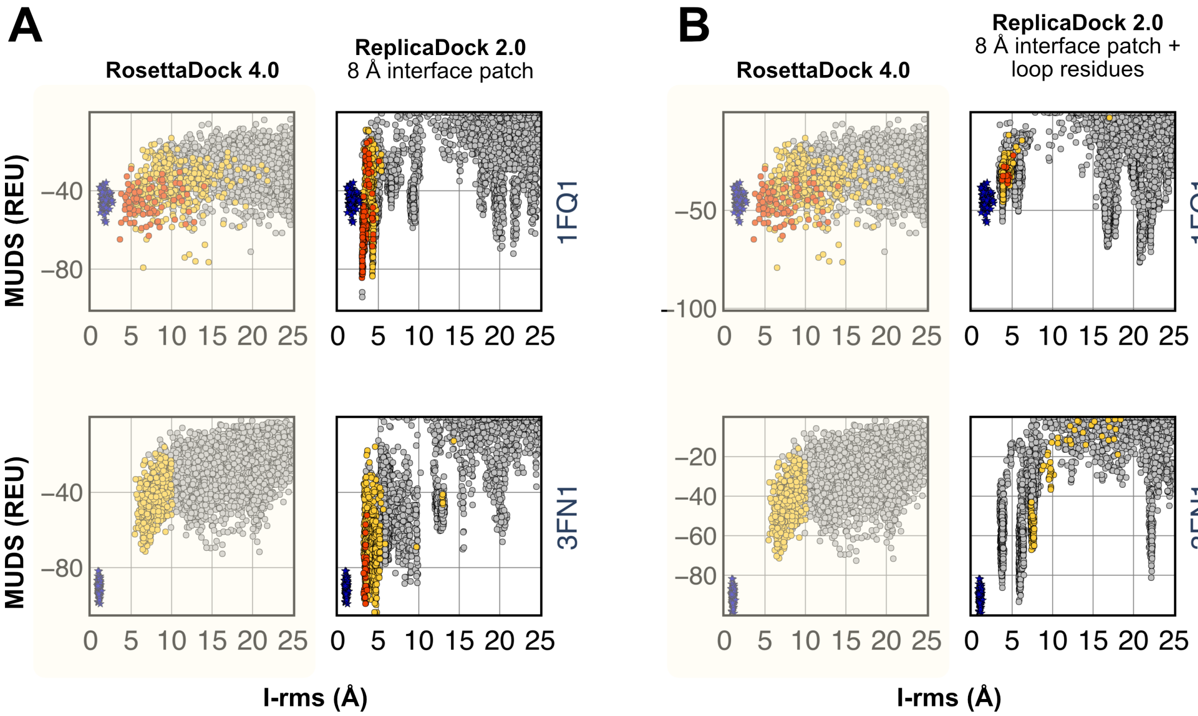


**Fig. S5.** MUDS versus C$\alpha$-RMSD(Å) plots for RosettaDock 4.0 and ReplicaDock 2.0 for two sets of residue selections in the low-resolution stage. Mobile residues sets are as follows: (left) 8 Å interface patch, and (right) 8 Å interface patch + loops. Candidate structures are colored by the CAPRI quality post all-atom refinement (colors : green = high quality, red = moderate quality, yellow = acceptable quality, gray = incorrect).

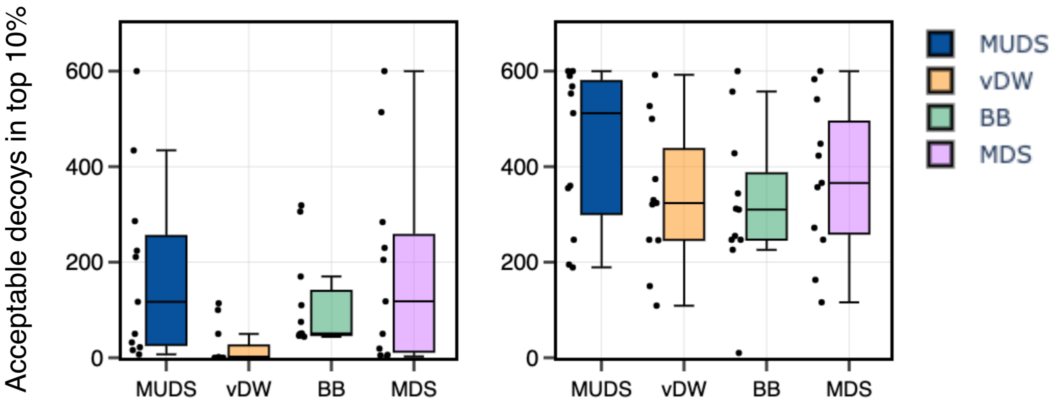


**Fig. S6. Near-native enrichment** for two different weights based on MUDS. Each score-term is represented with the boxplot illustrating the enrichment of CAPRI-quality accceptable models within the lowest-scoring 1,000 models out of a set of 10,000 (10%) for each of the 11 protein-protein complexes. The motif-dock score corresponds to the score function defined in *Marze et al.* (13). Motif Updated Dock Score(MUDS) *in blue* (as defined prior) comprises of Van der Waals score terms(attractive and repulsive clash terms) *in pale orange* and Backbone statistical terms(`rama`, `p_aa_pp` and `omega`) *in pale green* along with the motif-dock score *in purple*. (a) Represents reduced weights of 0.1X to the Van der Waals and the backbone score terms, where X is the standard weight as defined by `beta_nov16`. (b) Represents a weight of 1.0X to the score-terms where X is the standard weight defined by the `beta_nov16` score-function. Sampling performance is improved almost three-fold for all 11 targets by improved weights.

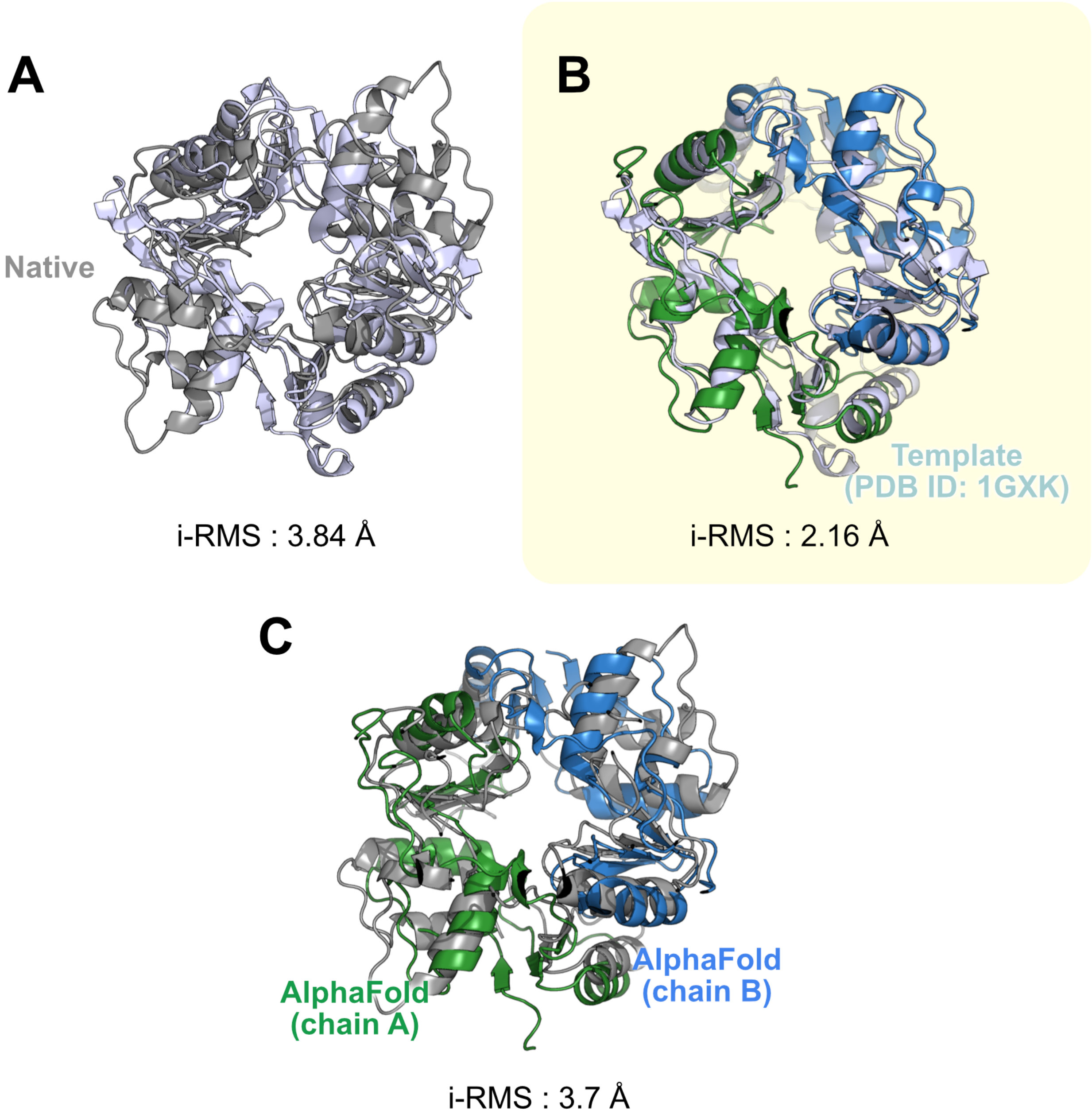


**Fig. S7.** AlphaFold complex modeling for CASP14-CAPRI target T164, comprising of the SMCHD1 (human) residues 1616-1899 Structural maintenance of chromosomes flexible hinge domain-containing protein 1. (A) Native structure (PDB ID: 6N64) in grey, superimposed by the available complex template (PDB ID: 1GXK) in light-blue, with an interface RMS of 3.84 Å. (B) AlphaFold model (green-blue) superimposed over the complex template (in light-blue). AlphaFold models the protein complex closer to the complex template, with considerable conformational differences amounting to an interface RMSD of 2.16 Å. (C) AlphaFold model superimposed over the native shows an interface RMSD of 3.7 Å. Although marked as an easy target due to the availability of the template, this target comprises flexible loops and helical rearrangements.

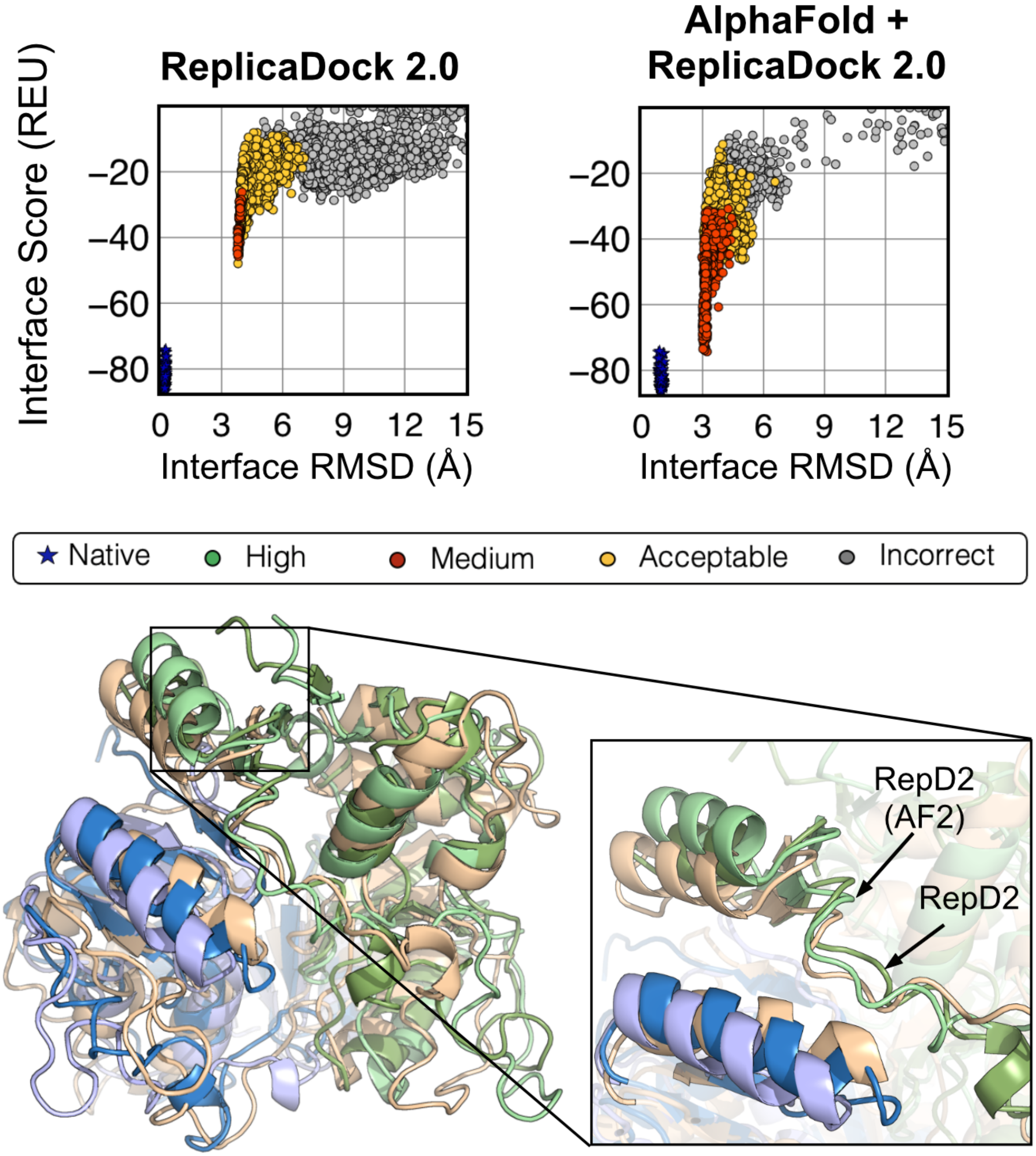


**Fig. S8.** ReplicaDock 2.0 performance on target T164 with and without AlphaFold template. (A) Interface Score (REU) vs Interface-RMSD (Å) plots show that ReplicaDock captures acceptable decoys (and a few medium-quality decoys) starting from the complex template, however, if flexible docking is initiated with AlphaFold structures, we improve the quality of our predictions by 0.6-0.8 Å and capture more medium-quality decoys with better interface scores. (B) The models from ReplicaDock with (green-blue) and without(palegreen-paleblue) AlphaFold template are superimposed over the native(grey-tan)

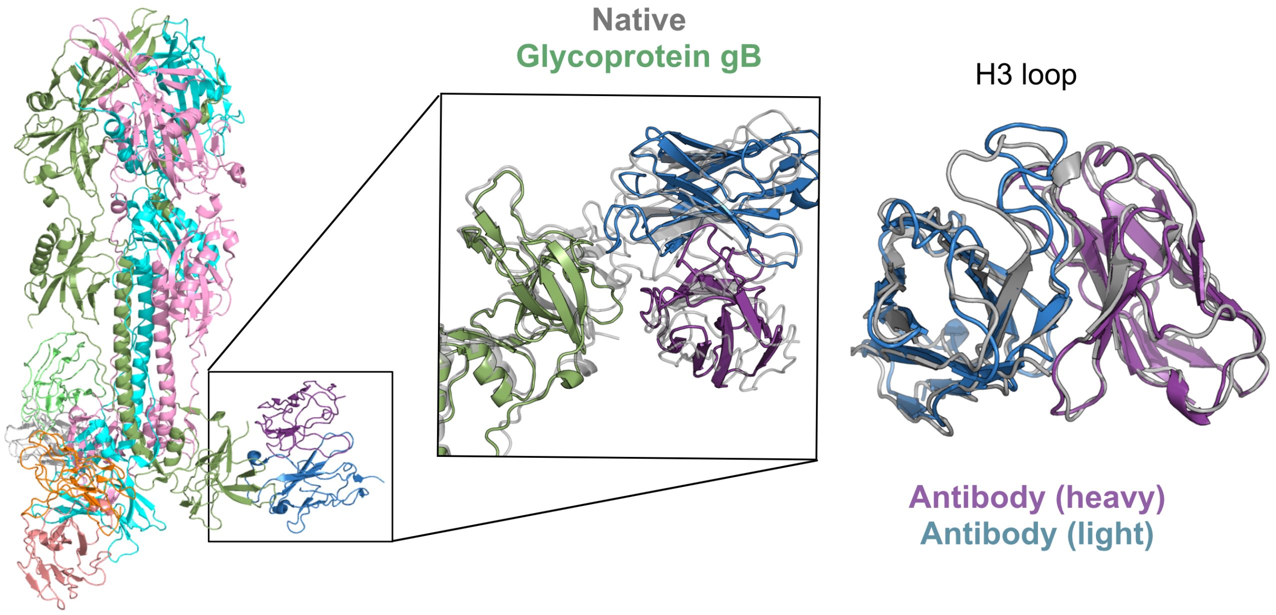


**Fig. S9.** ReplicaDock 2.0 performance on target T165 ( a monoclonal antibody 93k bound to varicella-zoster virus glycoprotein gB). AlphaFold predictions for complex predictions involved broken tertiary structures and are not reported. ReplicaDock found acceptable quality targets, however, the H3 loop was not adequately sampled owing to the relatively lower Interface-RMSD.

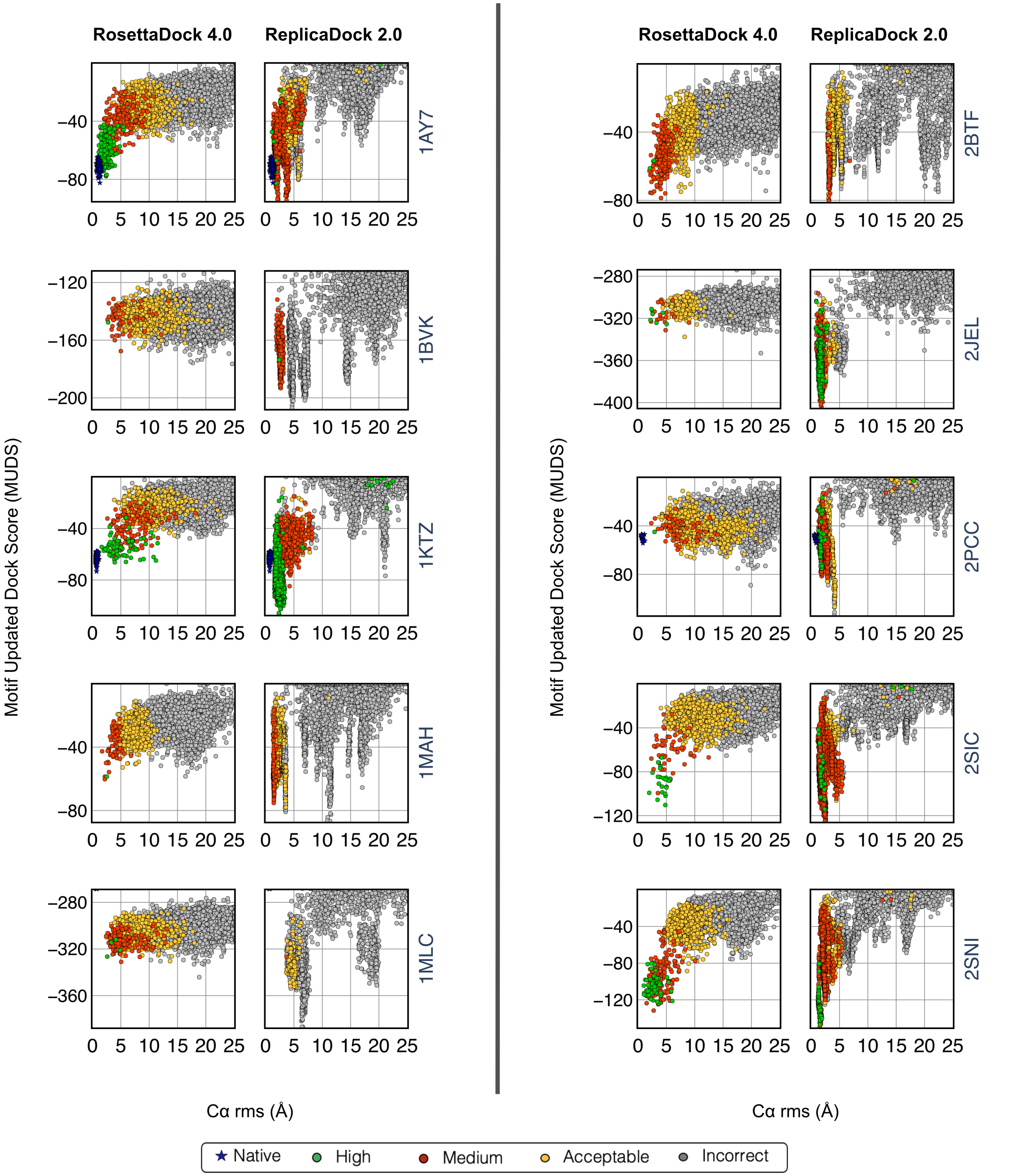


**Fig. S10.** Score versus Cα-RMSD(Å) plots in the low-resolution stage for motif updated dock score with RosettaDock 4.0 and ReplicaDock 2.0 for **rigid docking targets**.

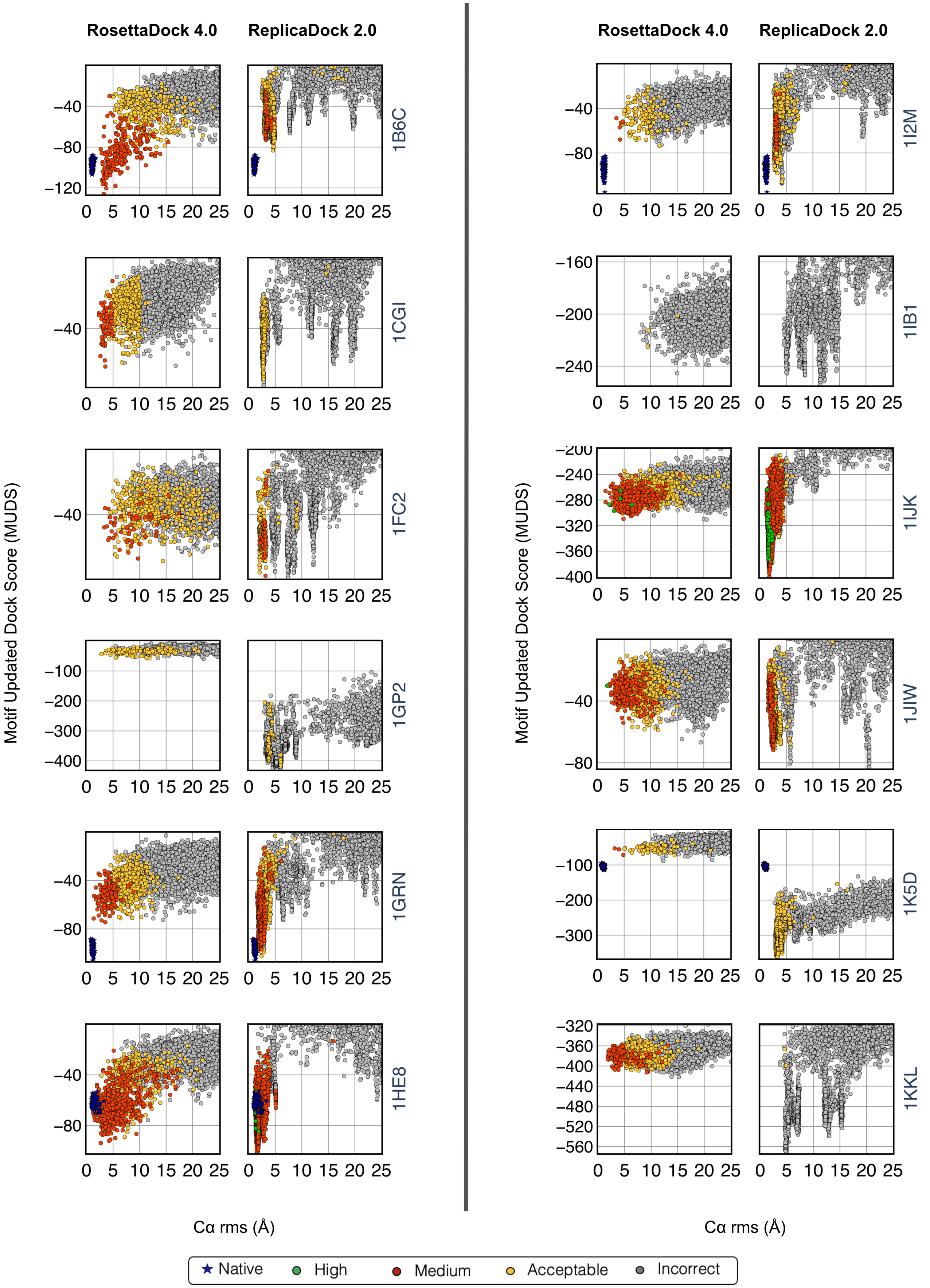
RosettaDock 4.0
ReplicaDock 2.0
1B6C
1CGI
1FC2
1GP2
1GRN
1HE8
1I2M
1IB1
1IJK
1JIW
1K5D
1KKL
Motif Updated Dock Score (MUDS)
Cα rms (Å)
Native
High
Medium
Acceptable
Incorrect

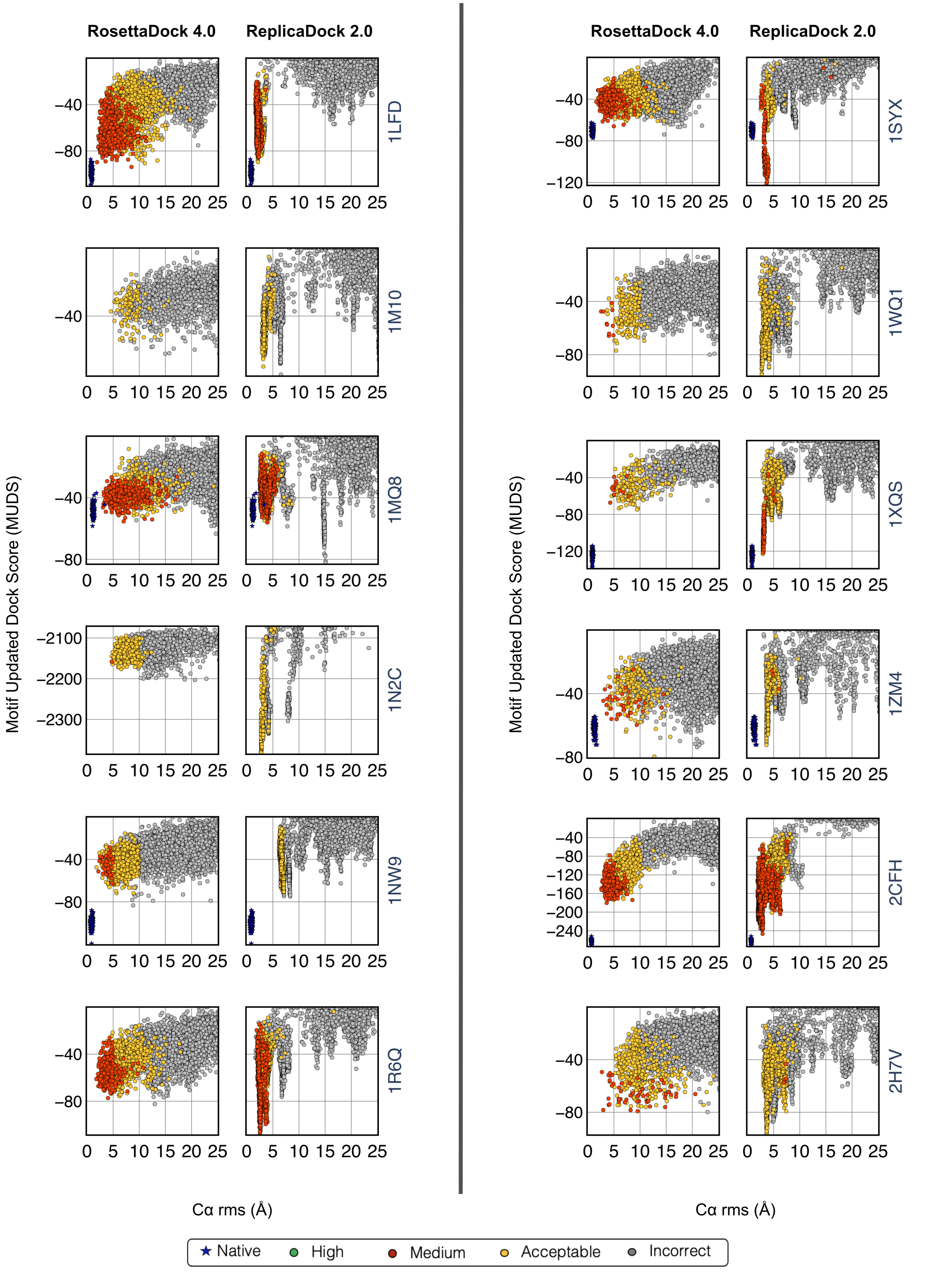
RosettaDock 4.0
ReplicaDock 2.0
1LFD
1M10
1MQ8
1N2C
1NW9
1R6Q
1SYX
1WQ1
1XQS
1ZM4
2CFH
2H7V
Motif Updated Dock Score (MUDS)
Cα rms (Å)
Native
High
Medium
Acceptable
Incorrect

 

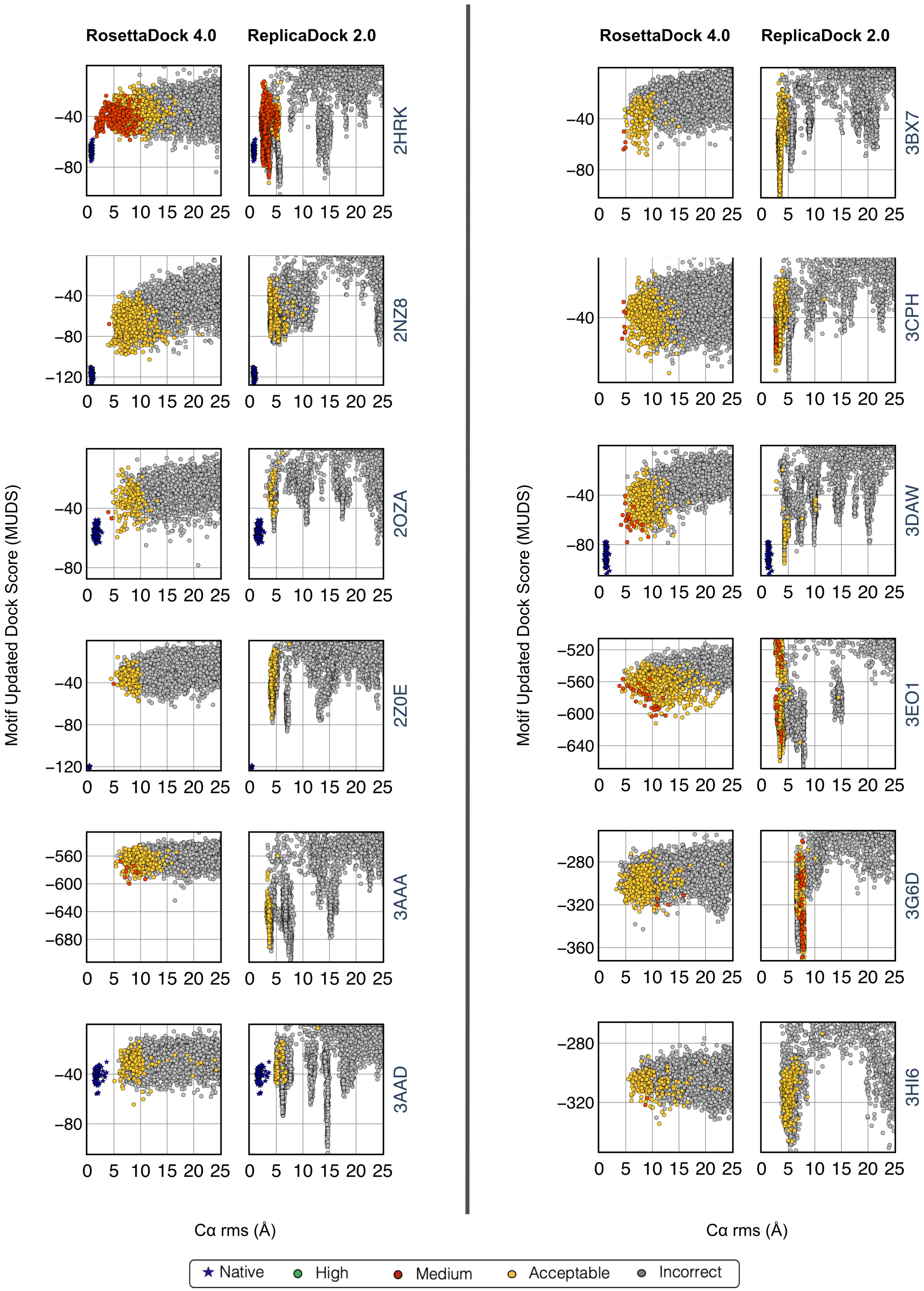

RosettaDock 4.0
ReplicaDock 2.0
2HRK
2NZ8
2OZA
2Z0E
3AAA
3AAD
3BX7
3CPH
3DAW
3EO1
3G6D
3HI6
Motif Updated Dock Score (MUDS)
Cα rms (Å)
Native
High
Medium
Acceptable
Incorrect

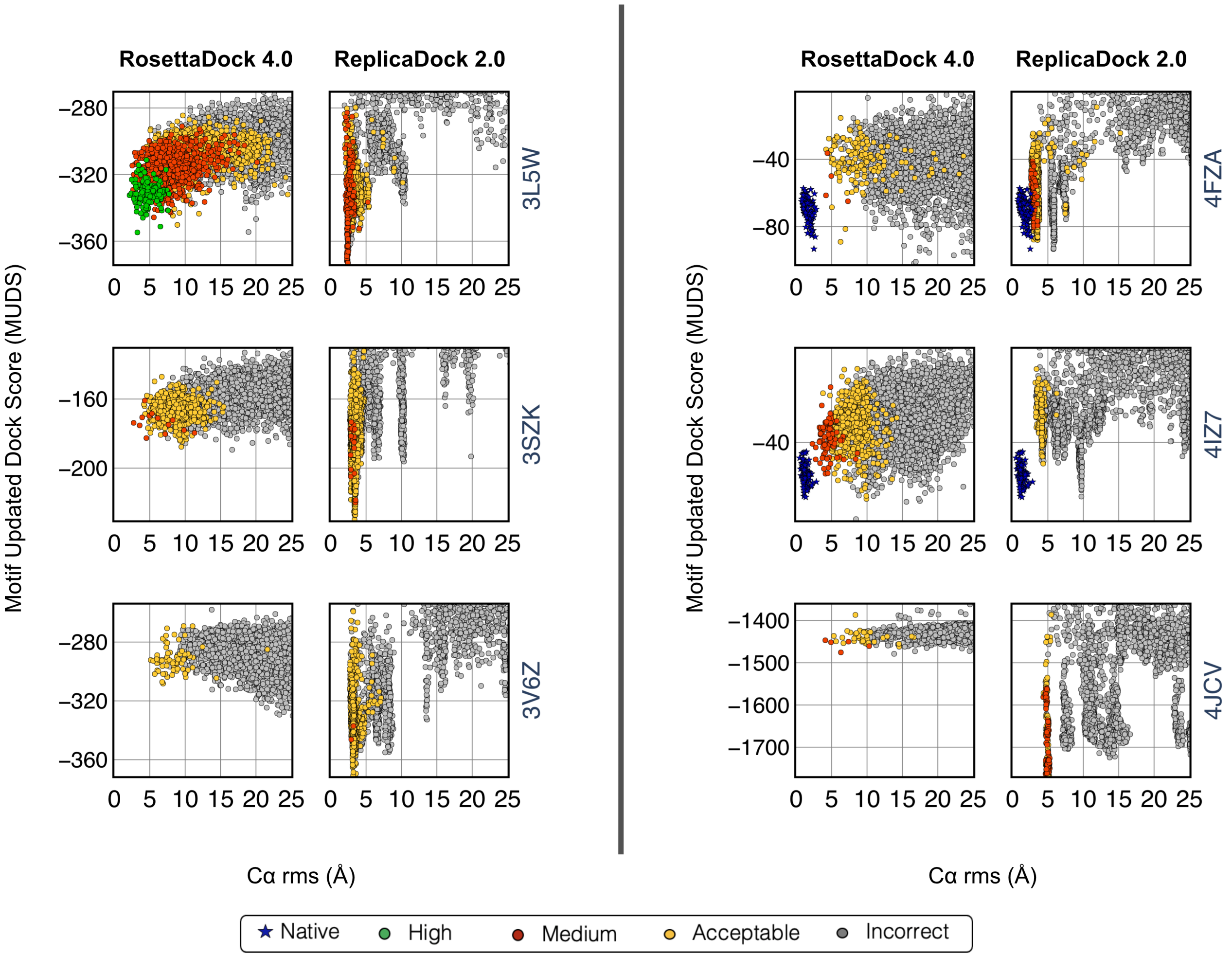


**Fig. S11.** Score versus C$\alpha$-RMSD(Å) plots in the low-resolution stage for motif updated dock score with RosettaDock 4.0 and ReplicaDock 2.0 for **medium docking targets.**

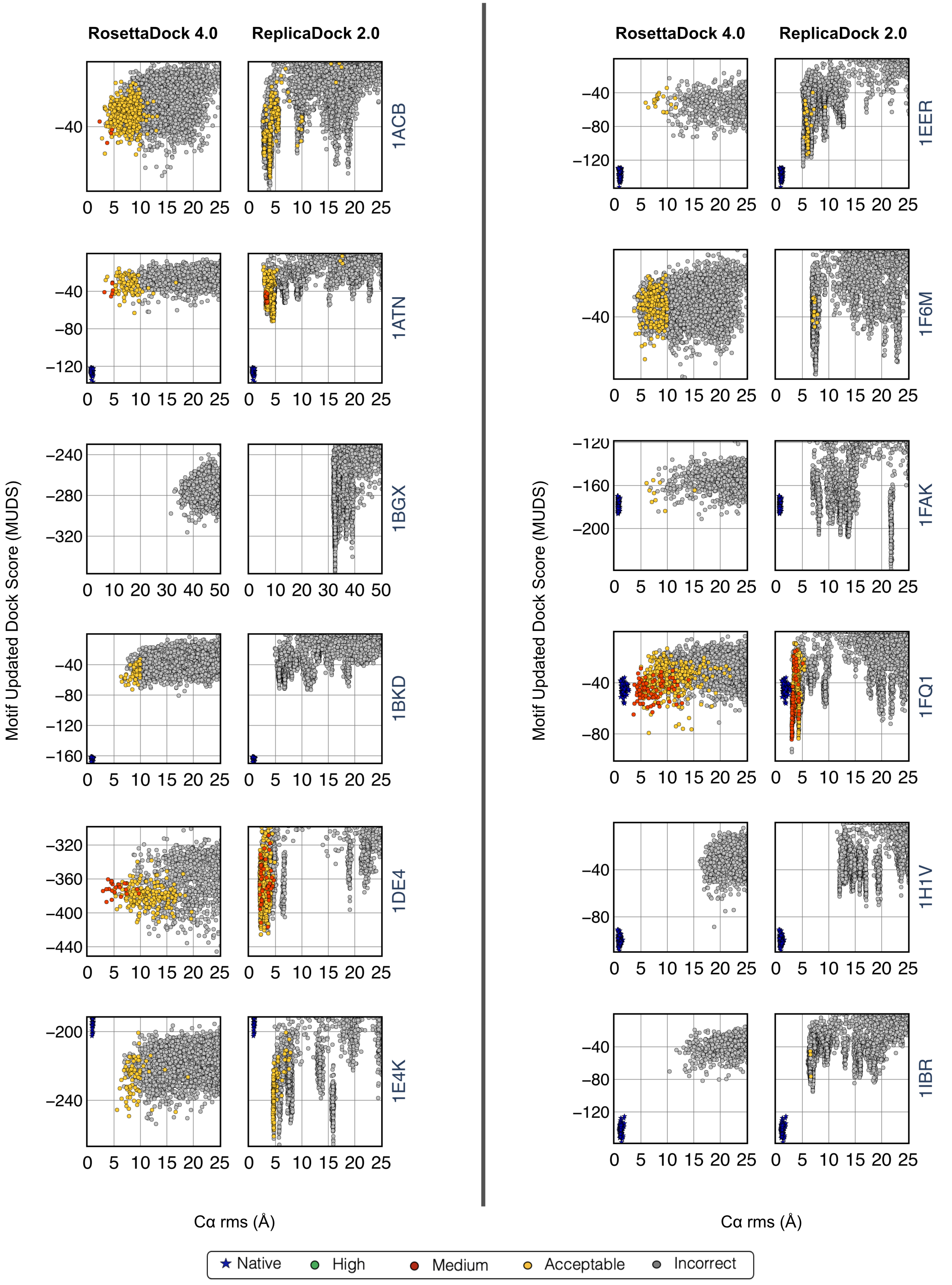
RosettaDock 4.0
ReplicaDock 2.0
1ACB
1ATN
1BGX
1BKD
1DE4
1E4K
1EER
1F6M
1FAK
1FQ1
1H1V
1IBR
Motif Updated Dock Score (MUDS)
Cα rms (Å)
Native
High
Medium
Acceptable
Incorrect

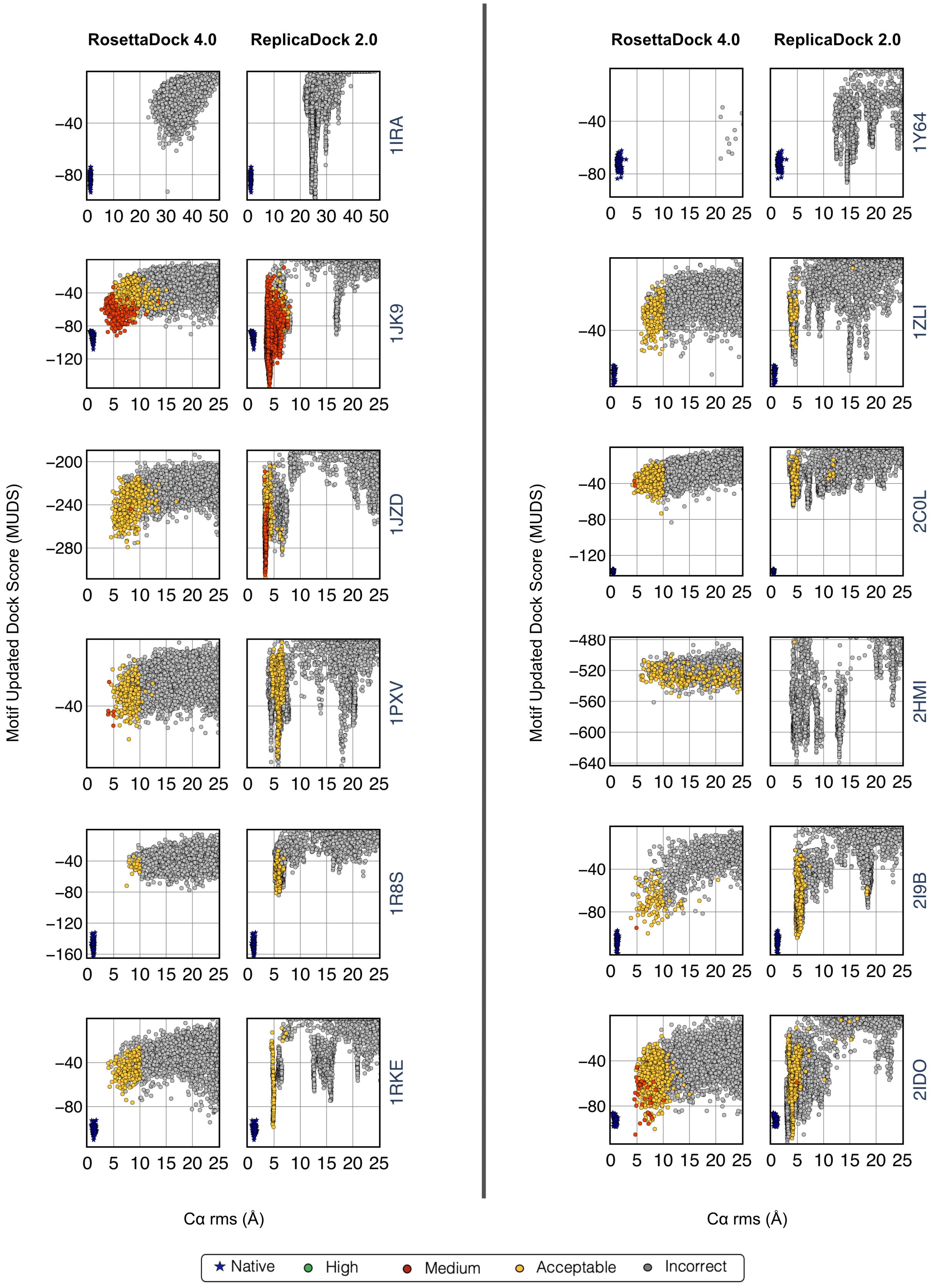
RosettaDock 4.0
ReplicaDock 2.0
RosettaDock 4.0
ReplicaDock 2.0
1IRA
1JK9
1JZD
1PXV
1R8S
1RKE
1Y64
1ZLI
2C0L
2HMI
2I9B
2IDO
Motif Updated Dock Score (MUDS)
Motif Updated Dock Score (MUDS)
Cα rms (Å)
Cα rms (Å)
Native
High
Medium
Acceptable
Incorrect

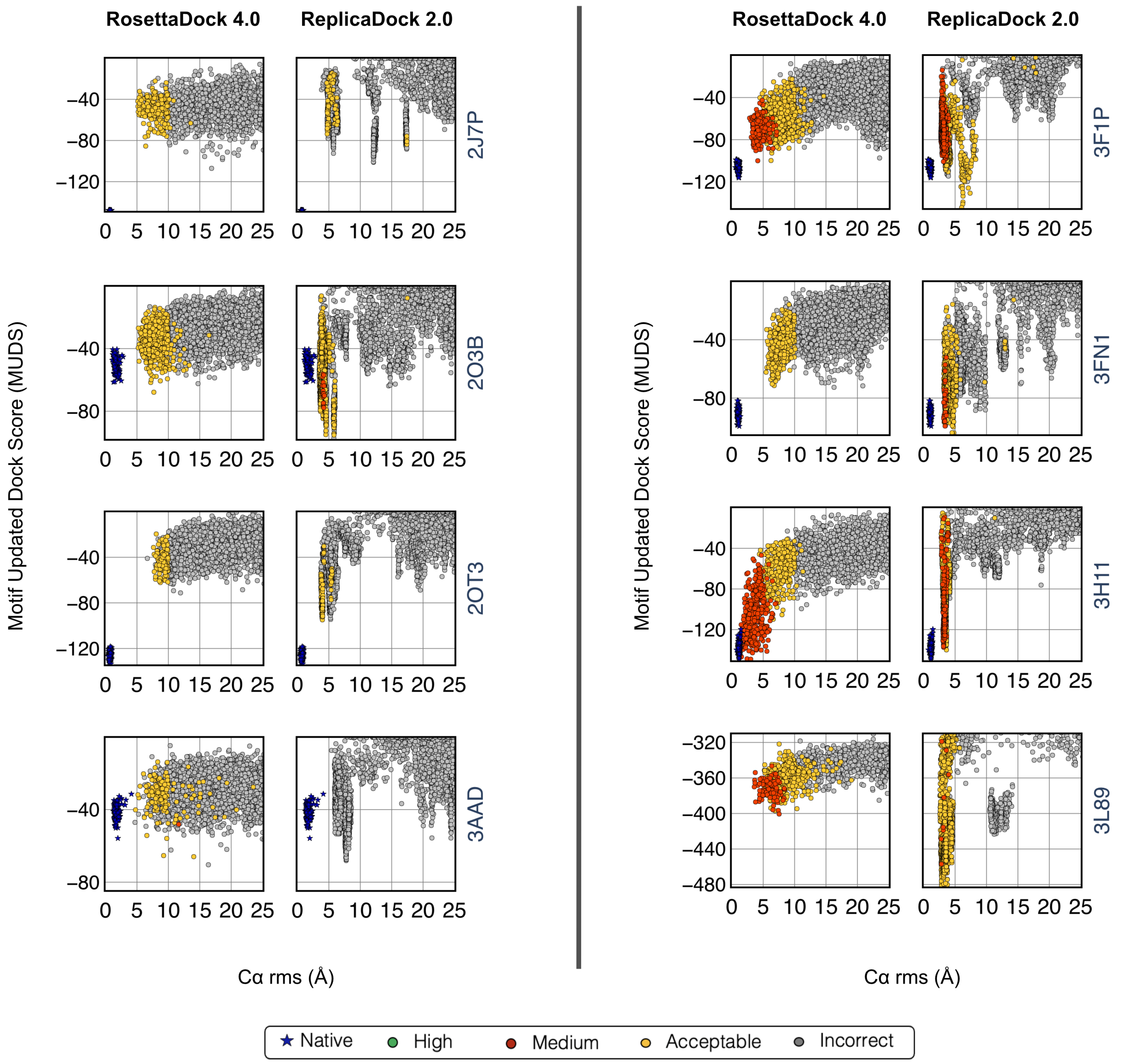


**Fig. S12.** Score versus C$\alpha$-RMSD(Å) plots in the low-resolution stage for motif updated dock score with RosettaDock 4.0 and ReplicaDock 2.0 for **difficult docking targets.**

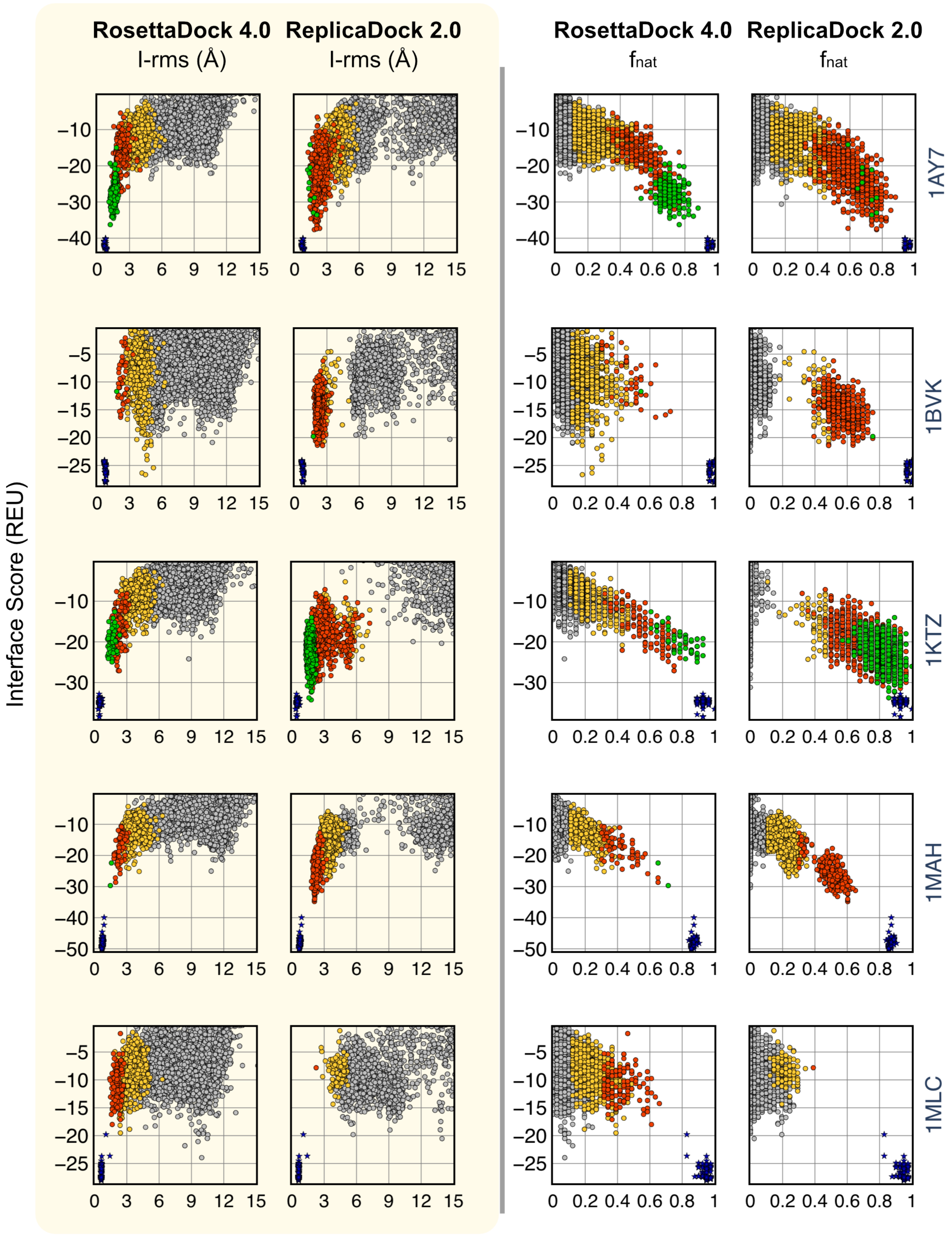

RosettaDock 4.0
ReplicaDock 2.0
I-rms (Å)
$f_{nat}$
Interface Score (REU)
1AY7
1BVK
1KTZ
1MAH
1MLC


 

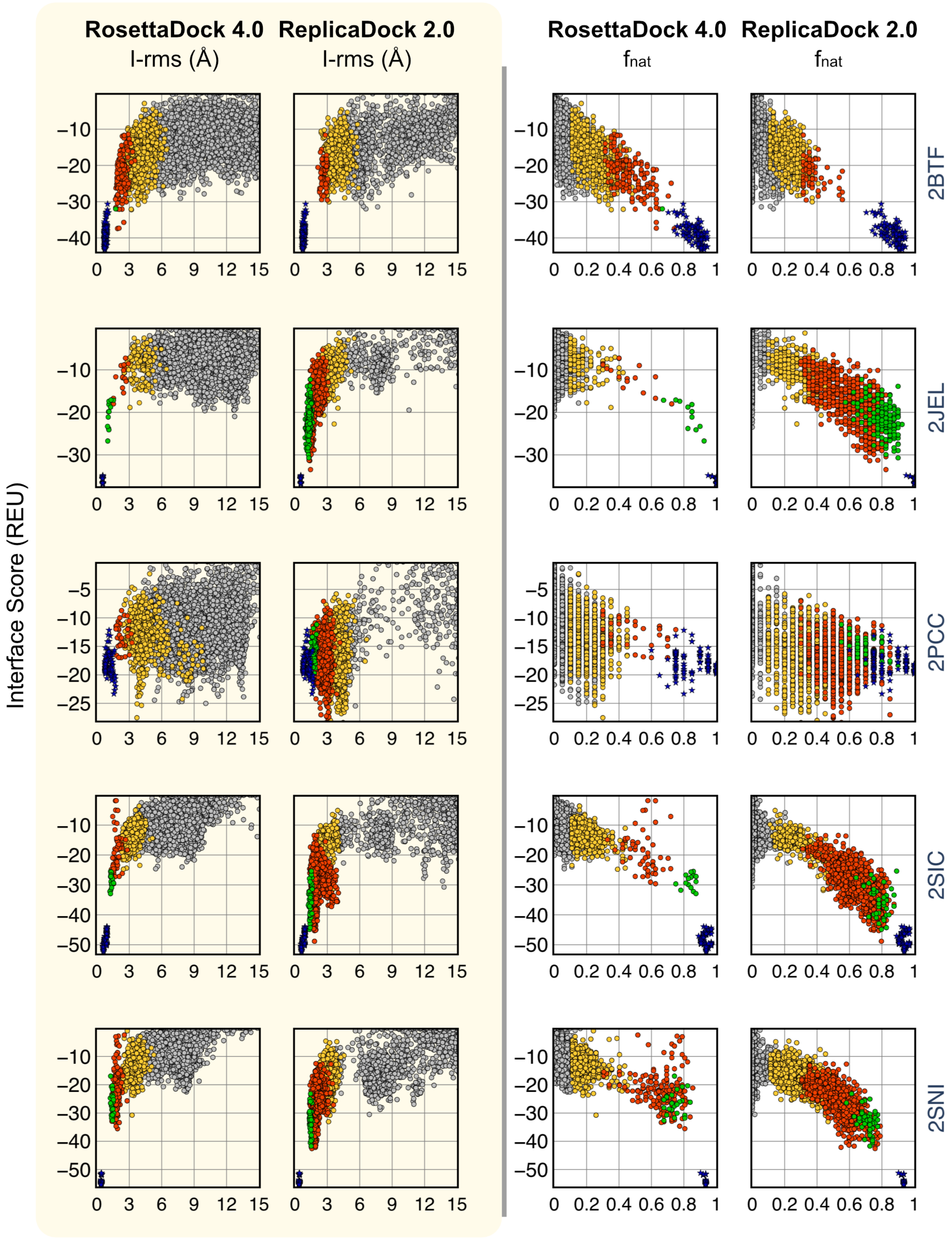


**Fig. S13.** Interface Score versus Interface-RMSD(Å) plots and Interface Score versus $f_{\mathrm{nat}}$ plots after the complete protocol for RosettaDock 4.0 and ReplicaDock 2.0 for rigid complexes.

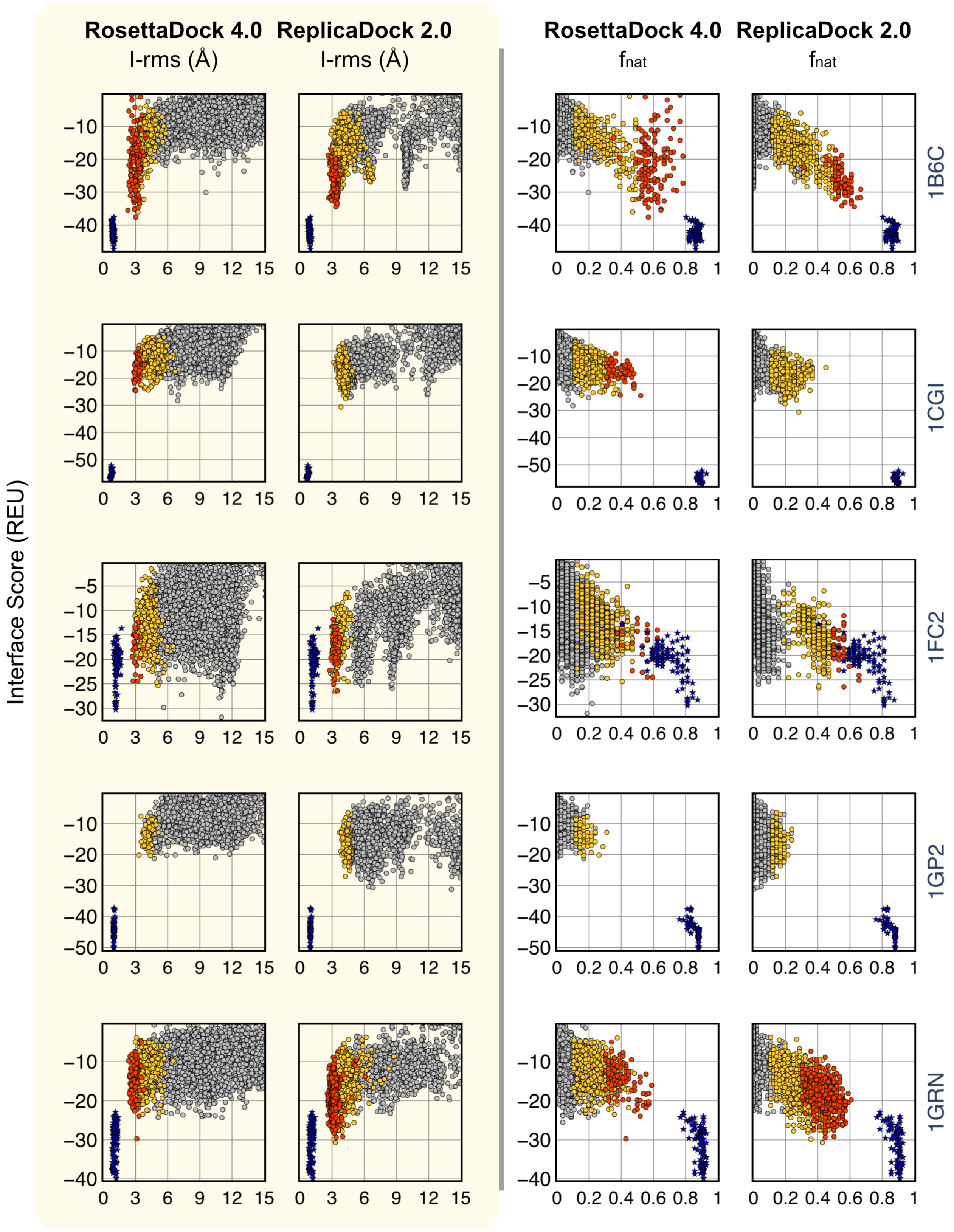



Ameya Harmalkar, Sai Pooja Mahajan and Jeffrey J. Gray

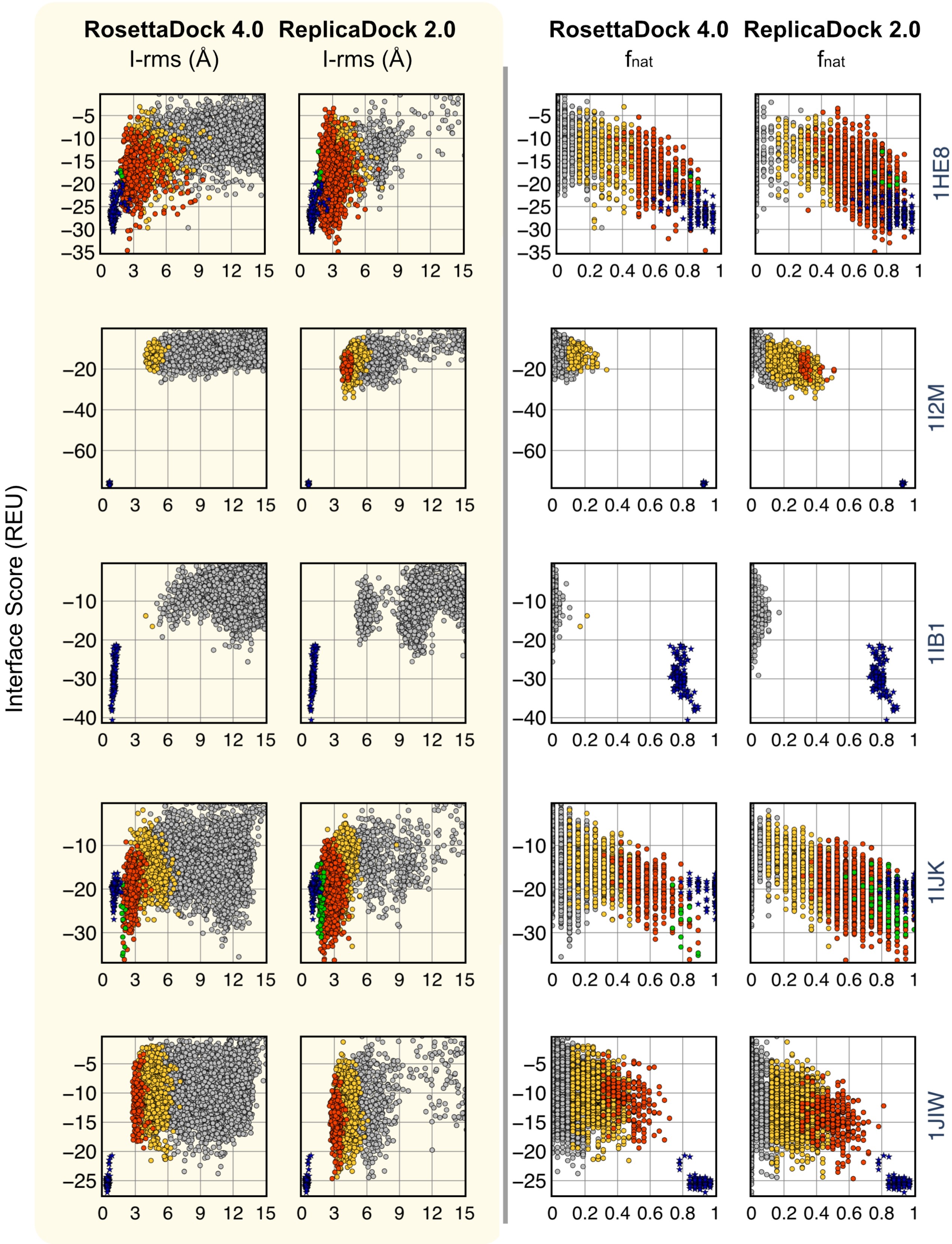
RosettaDock 4.0
ReplicaDock 2.0
I-rms (Å)
$f_{nat}$
Interface Score (REU)
1HE8
1I2M
1IB1
1IJK
1JIW

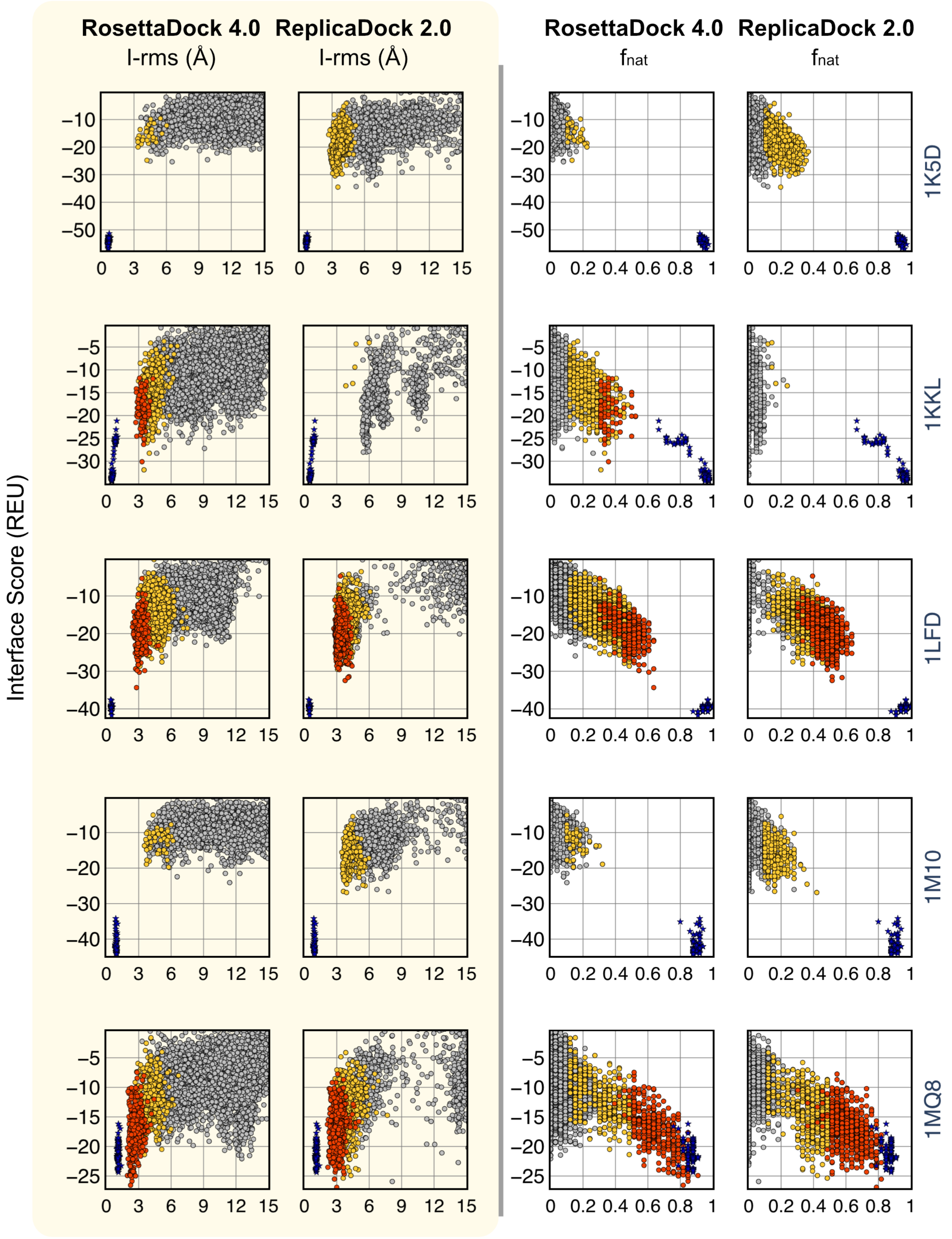
RosettaDock 4.0
I-rms (Å)
ReplicaDock 2.0
I-rms (Å)
RosettaDock 4.0
fnat
ReplicaDock 2.0
fnat
Interface Score (REU)
1K5D
1KKL
1LFD
1M10
1MQ8

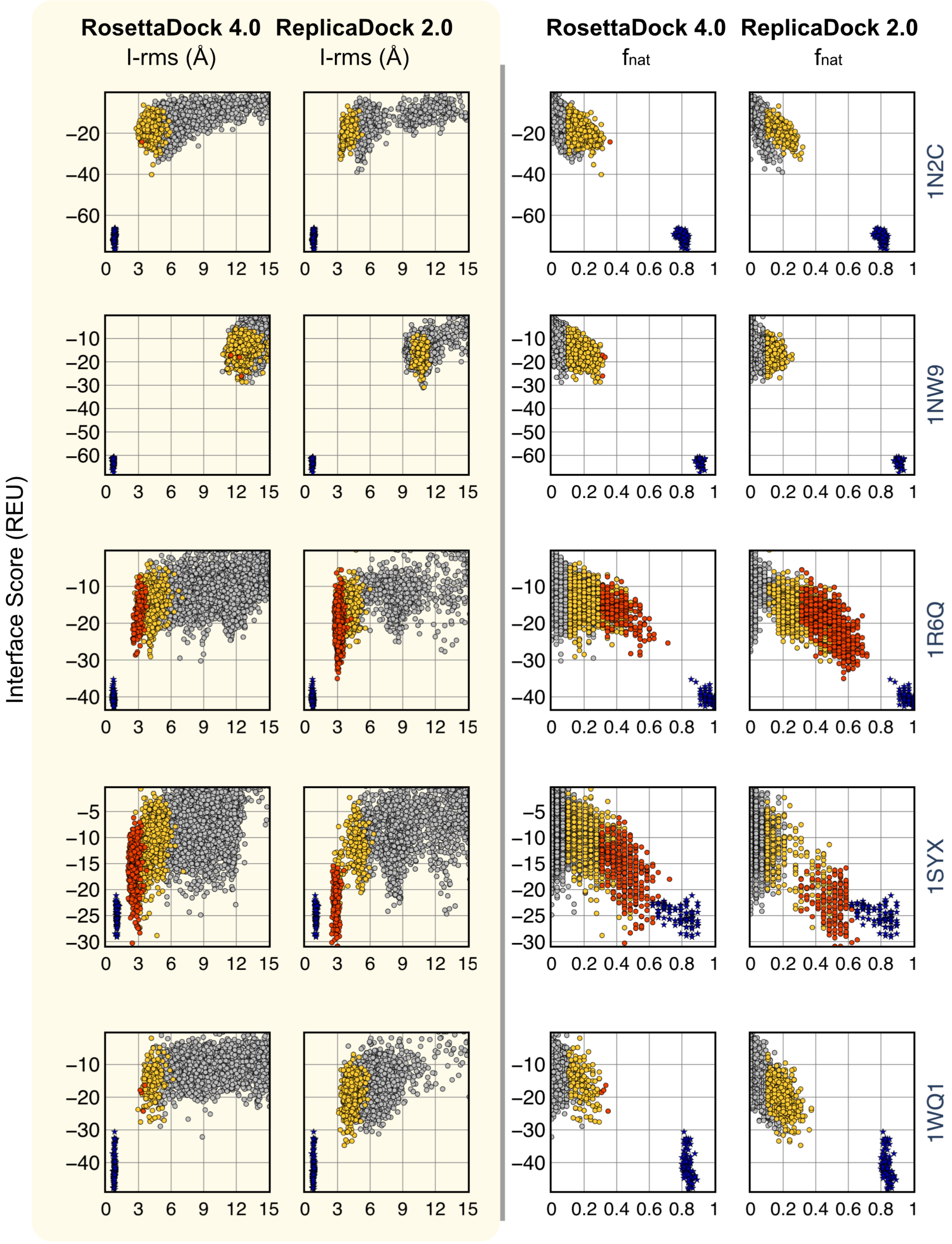
RosettaDock 4.0
ReplicaDock 2.0
I-rms (Å)
I-rms (Å)
RosettaDock 4.0
ReplicaDock 2.0
$f_{nat}$
$f_{nat}$
Interface Score (REU)
1N2C
1NW9
1R6Q
1SYX
1WQ1

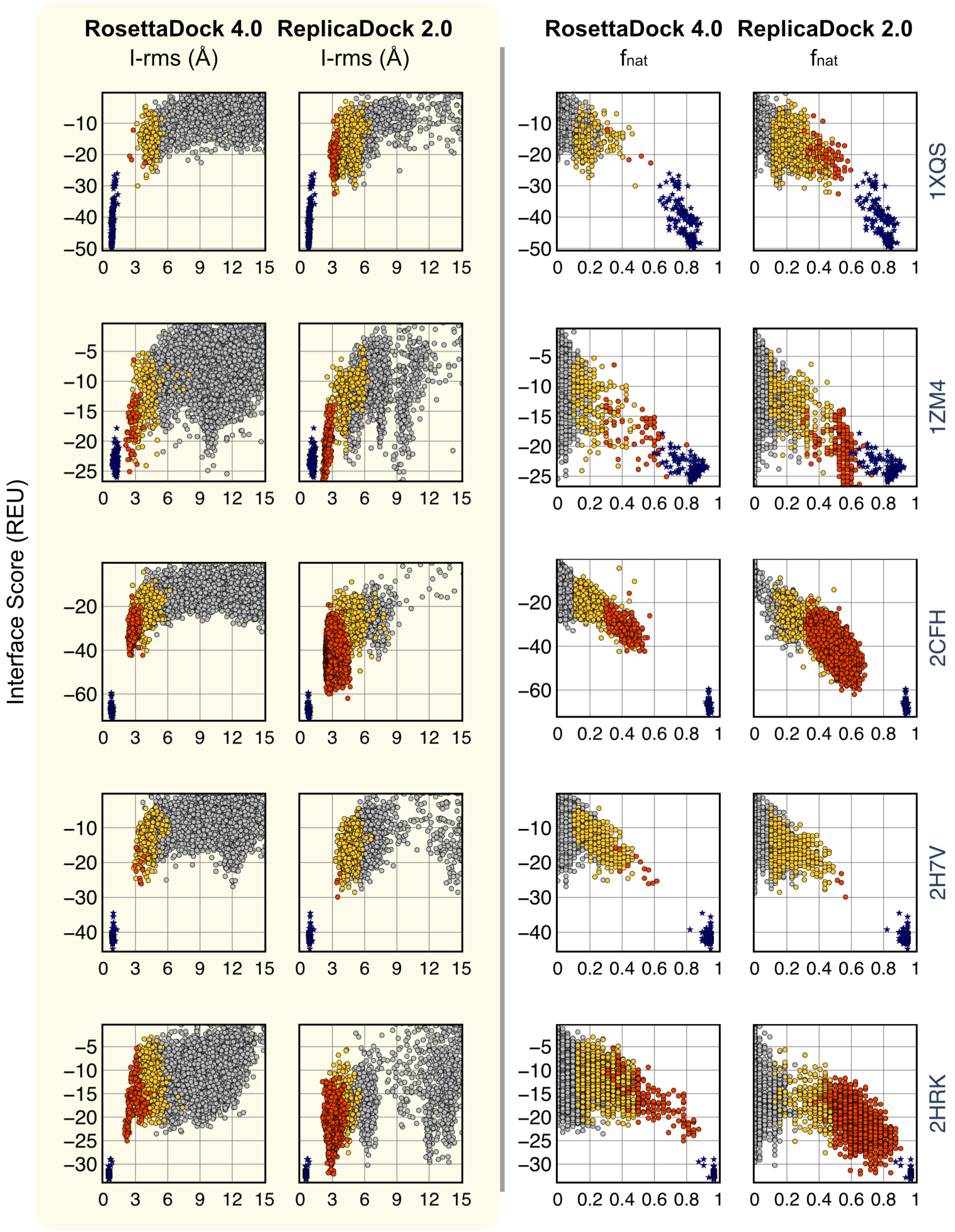
RosettaDock 4.0
ReplicaDock 2.0
I-rms (Å)
RosettaDock 4.0
ReplicaDock 2.0
$f_{nat}$
Interface Score (REU)
1XQS
1ZM4
2CFH
2H7V
2HRK

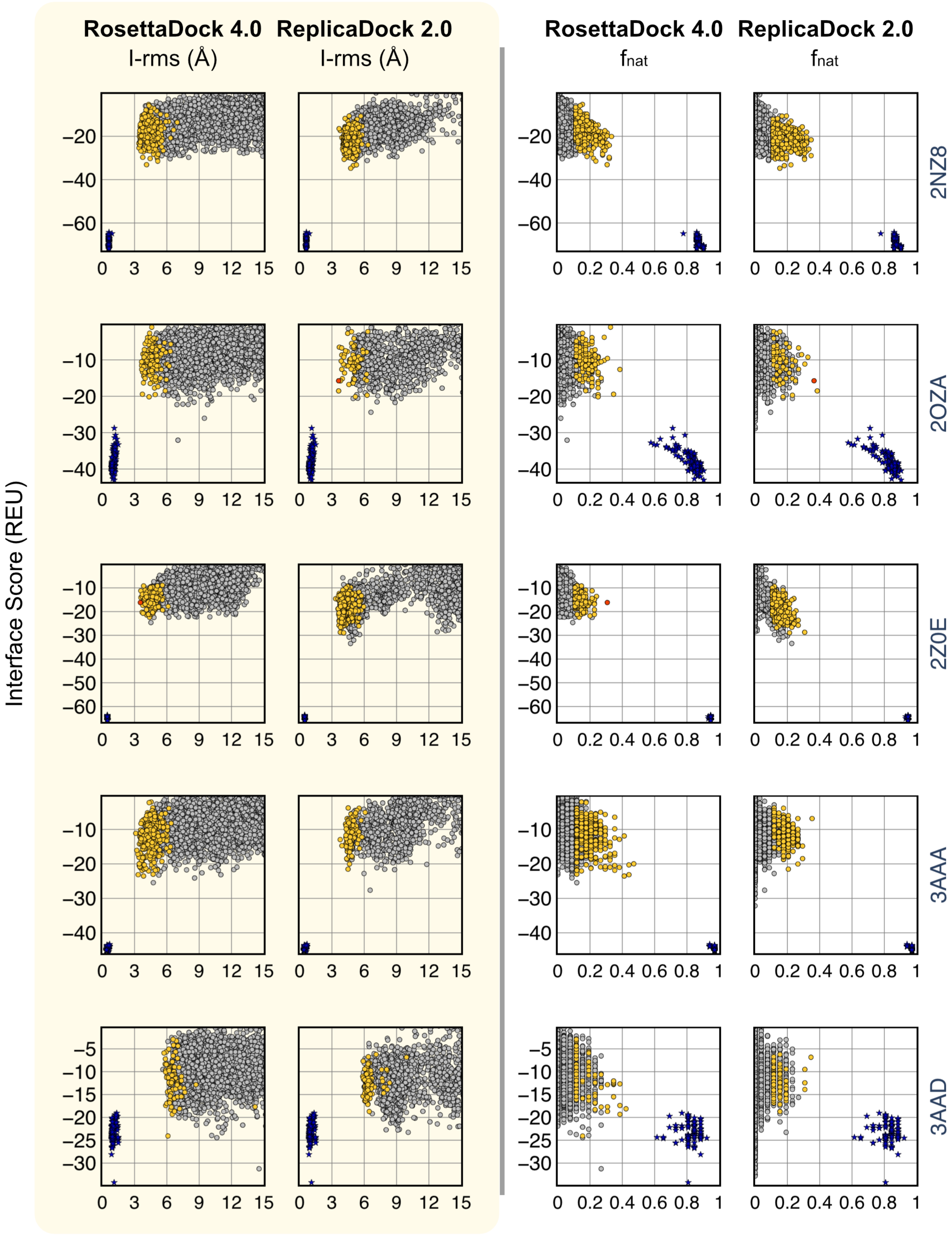
RosettaDock 4.0
ReplicaDock 2.0
I-rms (Å)
I-rms (Å)
RosettaDock 4.0
ReplicaDock 2.0
fnat
fnat
Interface Score (REU)
2NZ8
2OZA
2Z0E
3AAA
3AAD

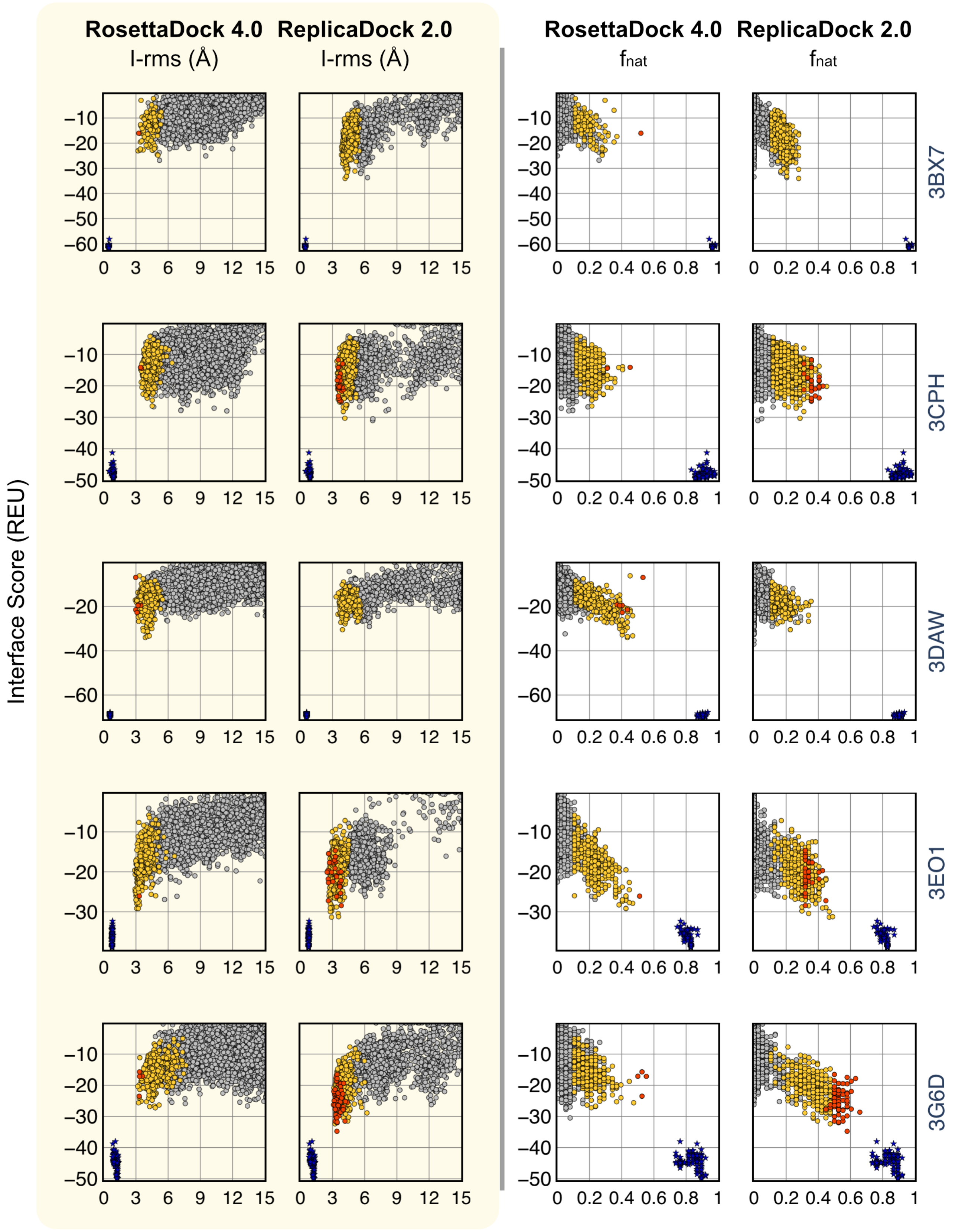
RosettaDock 4.0
I-rms (Å)
ReplicaDock 2.0
I-rms (Å)
RosettaDock 4.0
fnat
ReplicaDock 2.0
fnat
Interface Score (REU)
3BX7
3CPH
3DAW
3EO1
3G6D

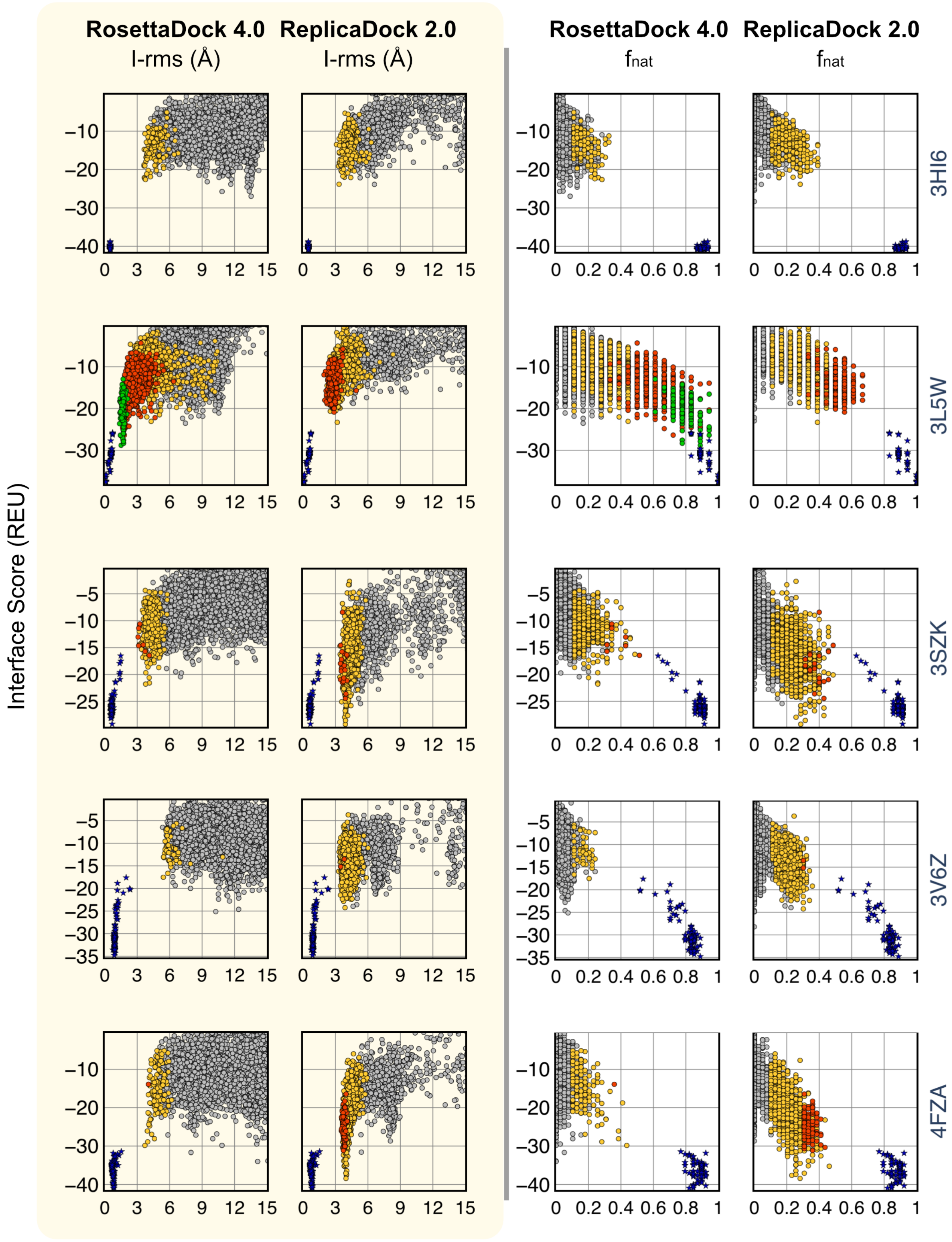

RosettaDock 4.0
I-rms (Å)
ReplicaDock 2.0
I-rms (Å)
RosettaDock 4.0
fnat
ReplicaDock 2.0
fnat
Interface Score (REU)
3HI6
3L5W
3SZK
3V6Z
4FZA

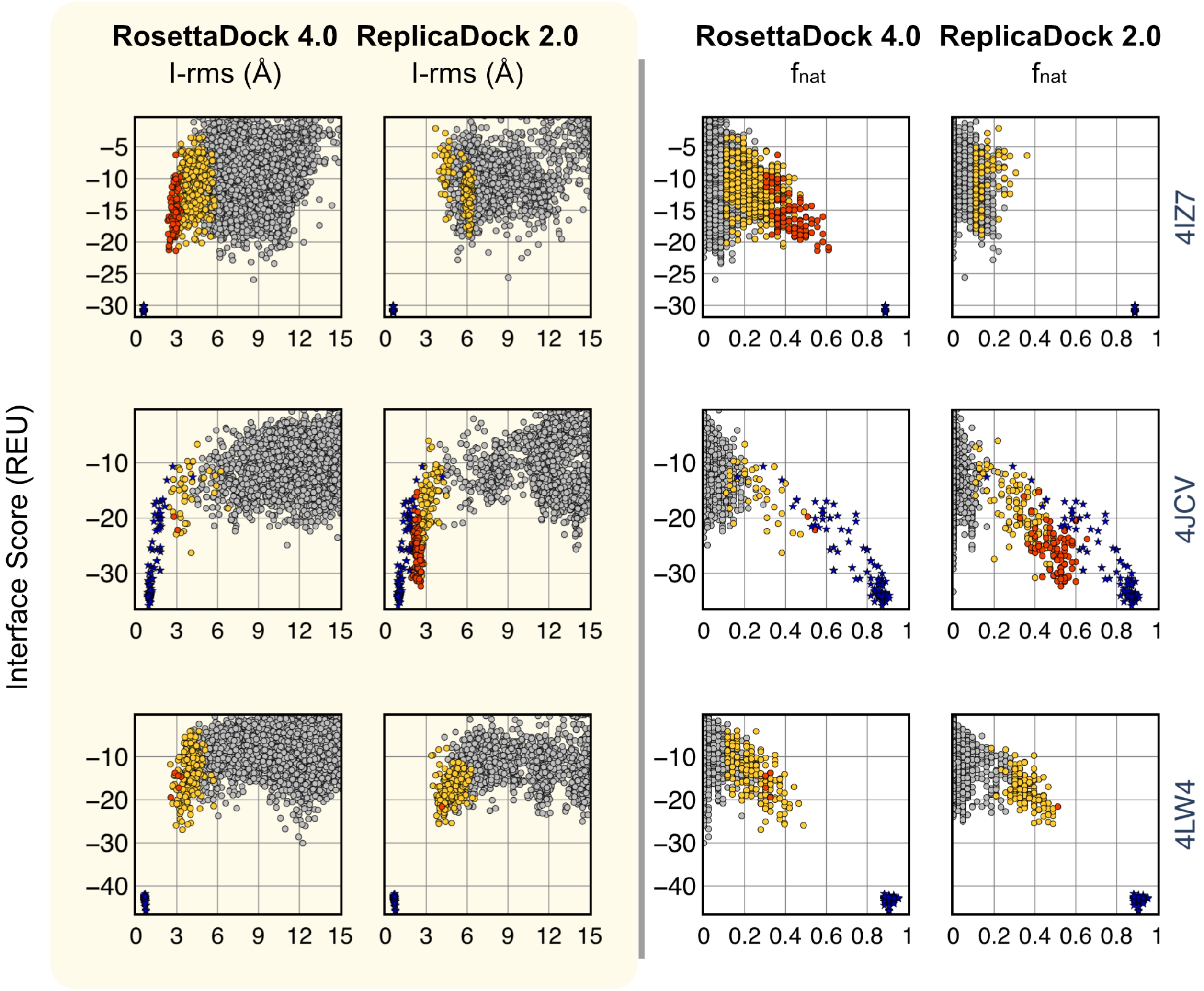


**Fig. S14.** Interface Score versus Interface-RMSD(Å) plots and Interface Score versus $f_{\mathrm{nat}}$ plots after the complete protocol for RosettaDock 4.0 and ReplicaDock 2.0 for **medium docking targets**.

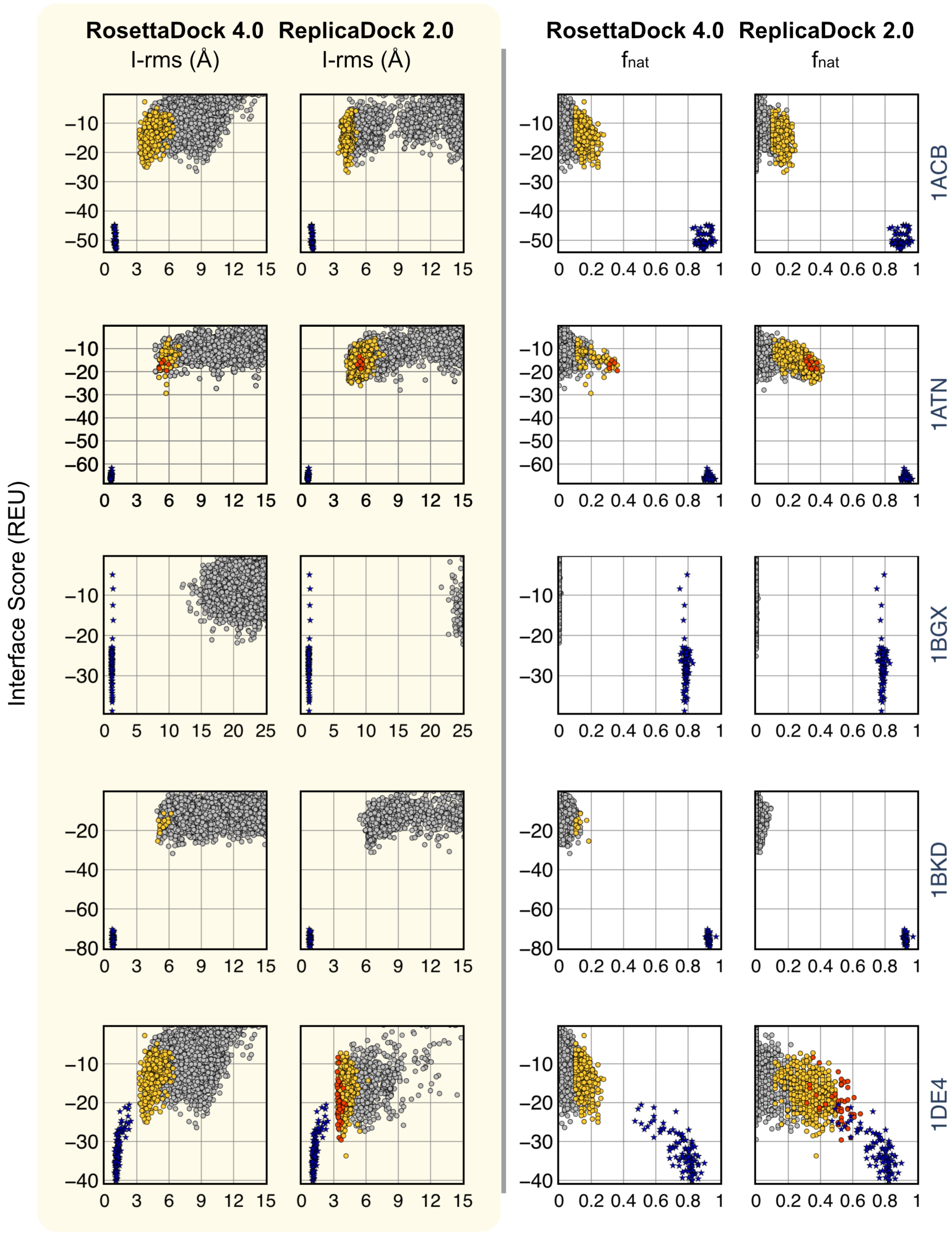

RosettaDock 4.0
ReplicaDock 2.0
I-rms (Å)
fnat
Interface Score (REU)
1ACB
1ATN
1BGX
1BKD
1DE4

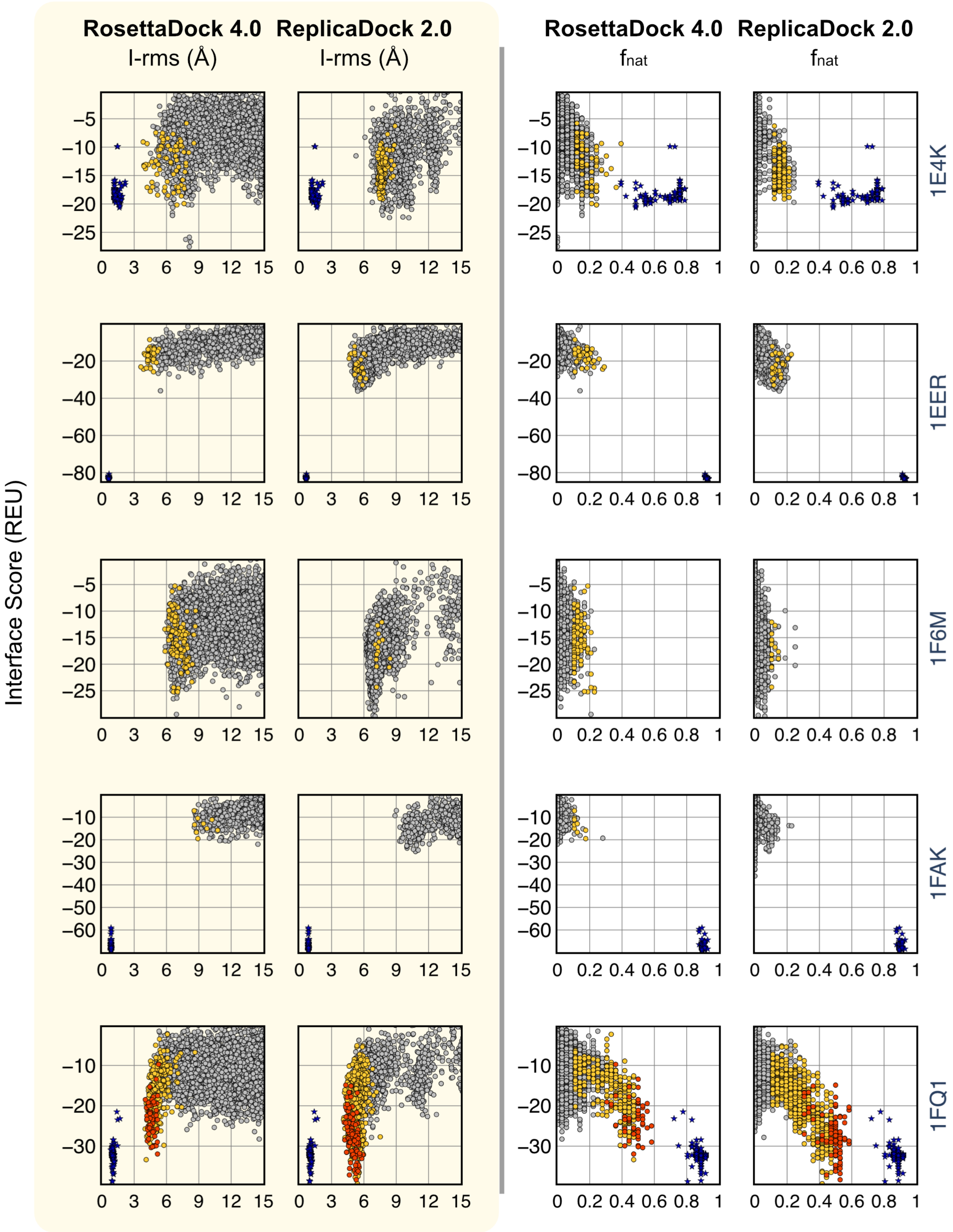
RosettaDock 4.0
ReplicaDock 2.0
I-rms (Å)
I-rms (Å)
RosettaDock 4.0
ReplicaDock 2.0
fnat
fnat
Interface Score (REU)
1E4K
1EER
1F6M
1FAK
1FQ1

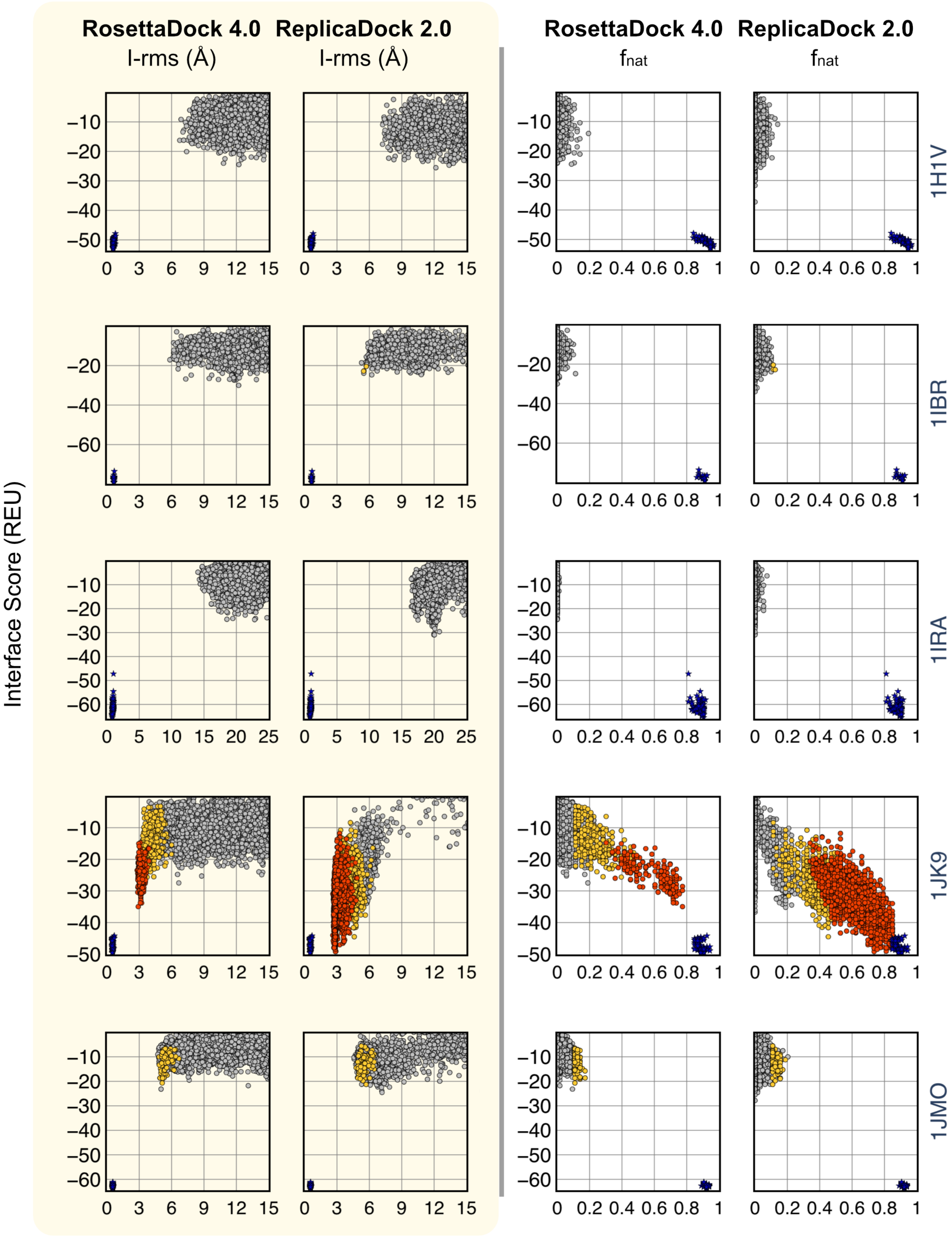
RosettaDock 4.0
ReplicaDock 2.0
I-rms (Å)
fnat
Interface Score (REU)
1H1V
1IBR
1IRA
1JK9
1JMO

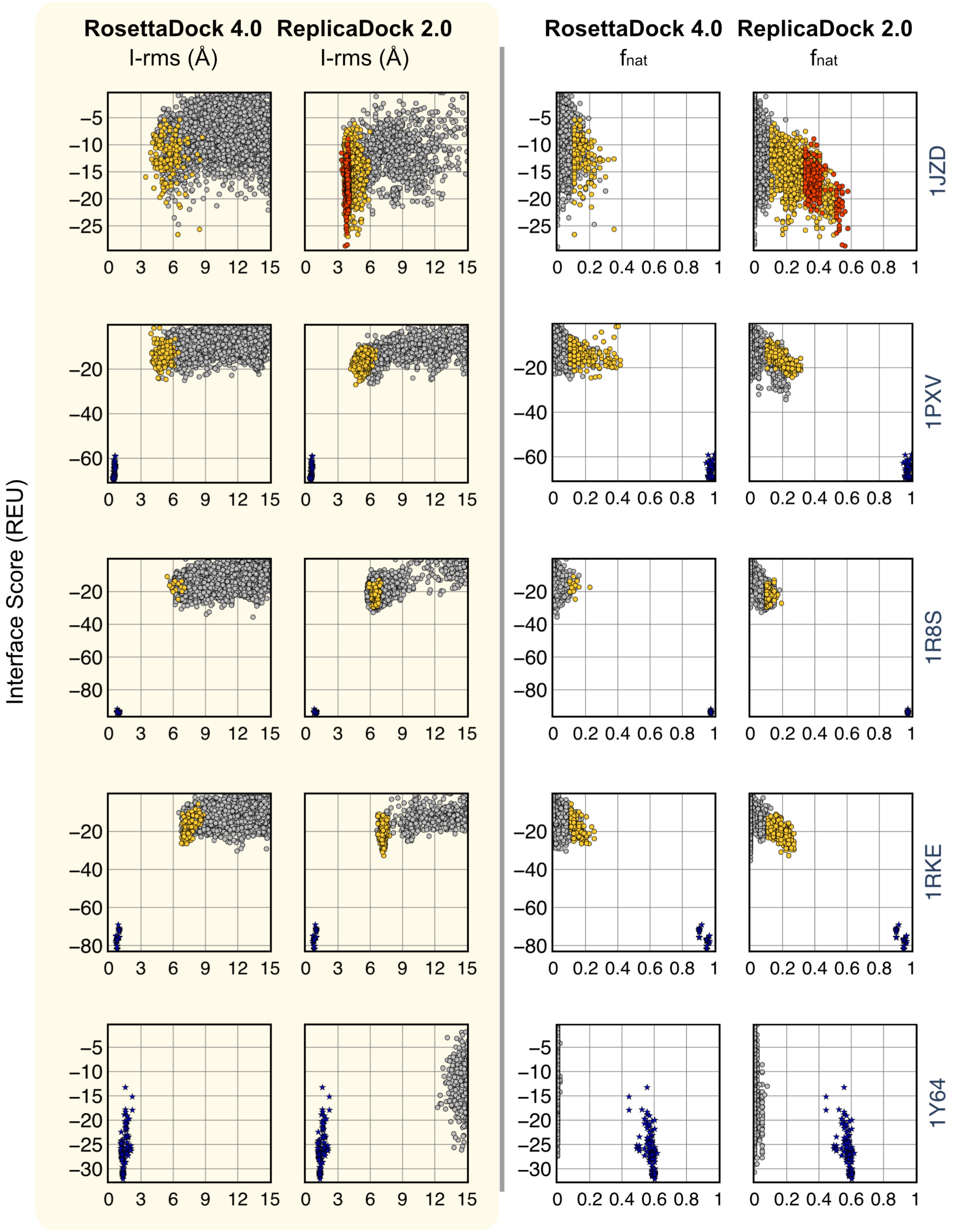

RosettaDock 4.0
ReplicaDock 2.0
I-rms (Å)
fnat
Interface Score (REU)
1JZD
1PXV
1R8S
1RKE
1Y64

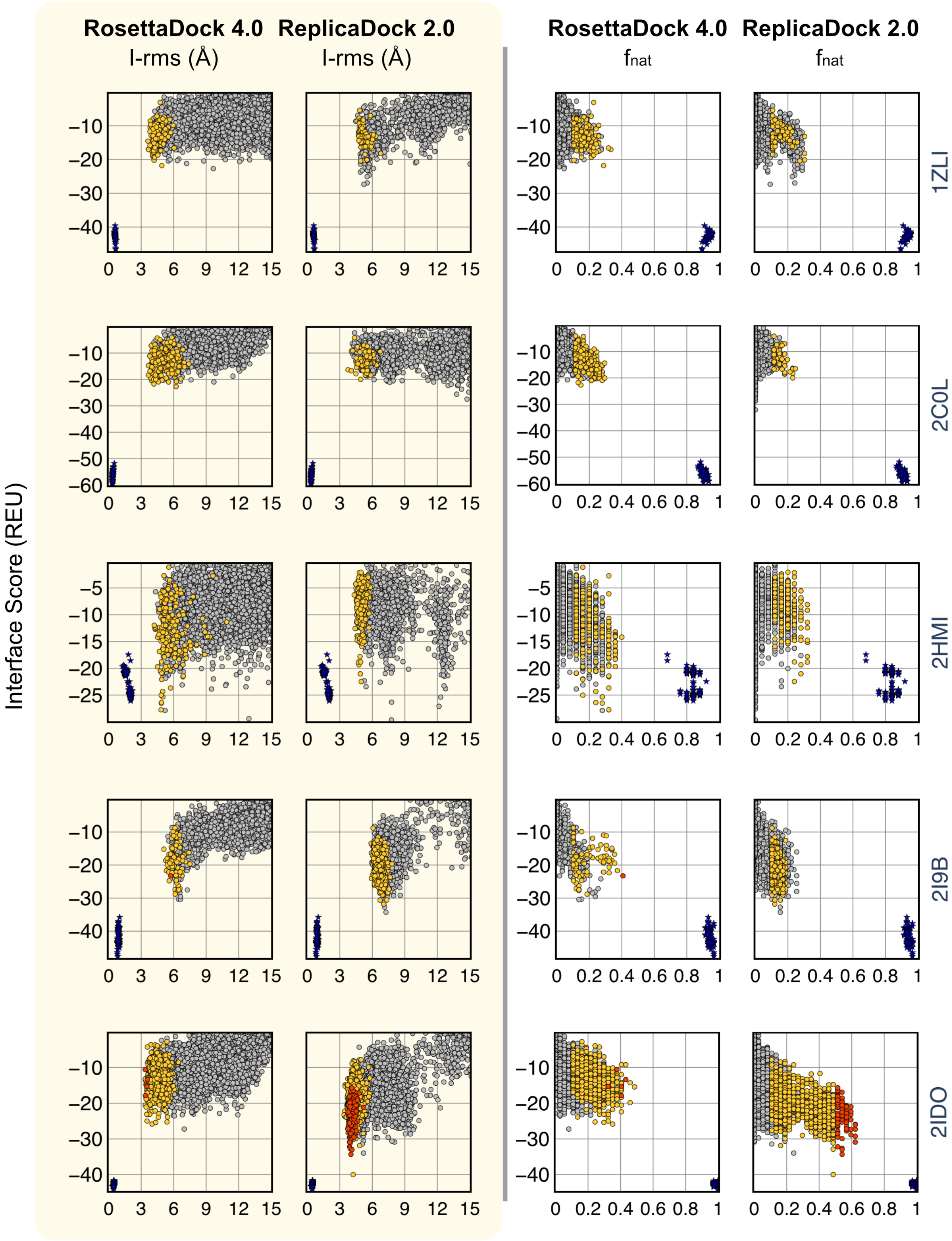
RosettaDock 4.0
ReplicaDock 2.0
I-rms (Å)
I-rms (Å)
RosettaDock 4.0
ReplicaDock 2.0
fnat
fnat
Interface Score (REU)
1ZLI
2C0L
2HMI
2I9B
2IDO

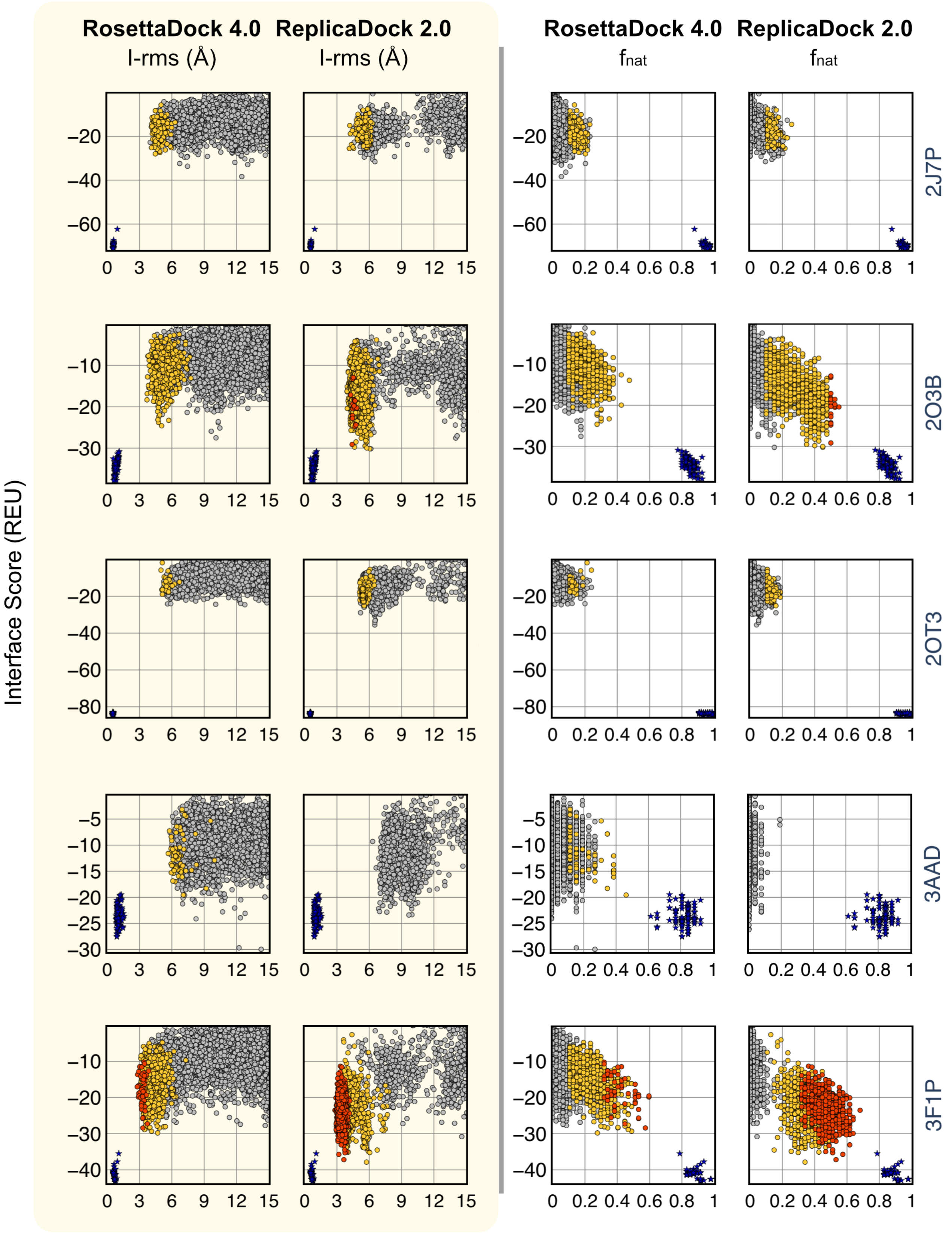
RosettaDock 4.0
ReplicaDock 2.0
I-rms (Å)
f$_{nat}$
Interface Score (REU)
2J7P
2O3B
2OT3
3AAD
3F1P

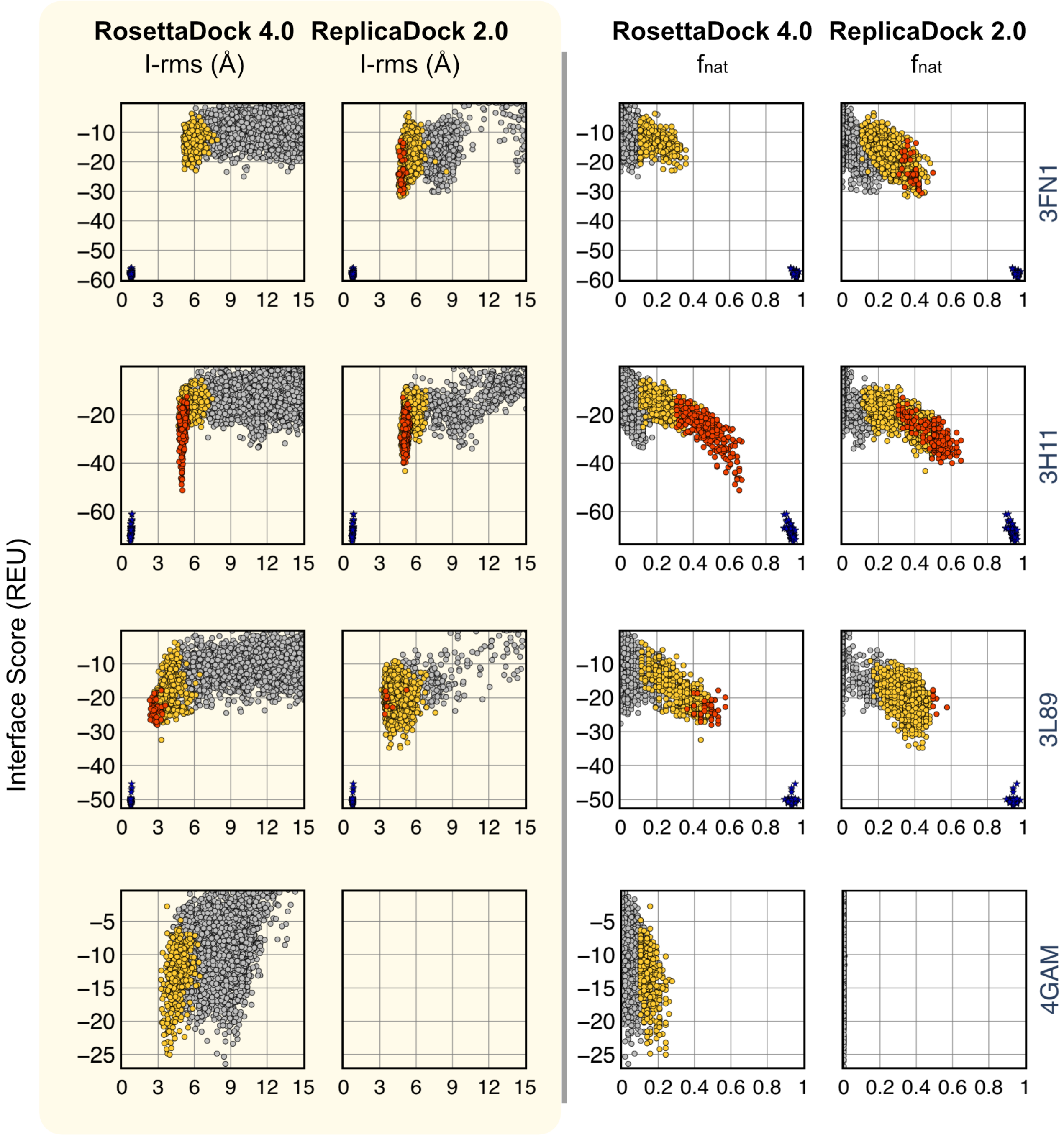


**Fig. S15.** Interface Score versus Interface-RMSD(Å) plots and Interface Score versus $f_{\mathrm{nat}}$ plots after the complete protocol for RosettaDock 4.0 and ReplicaDock 2.0 for **difficult docking targets**.

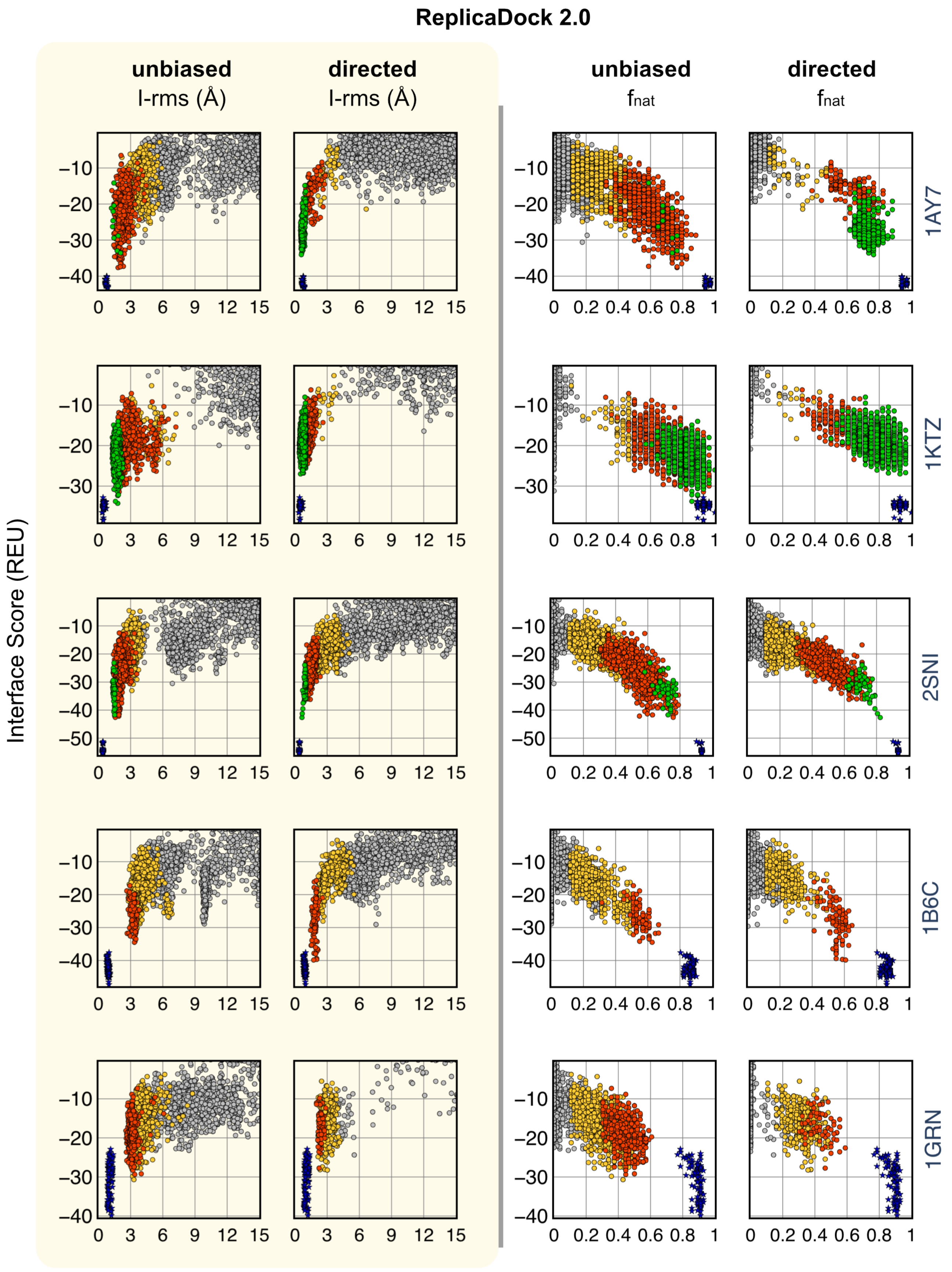
ReplicaDock 2.0
unbiased
I-rms (Å)
directed
I-rms (Å)
unbiased
fnat
directed
fnat
Interface Score (REU)
1AY7
1KTZ
2SNI
1B6C
1GRN

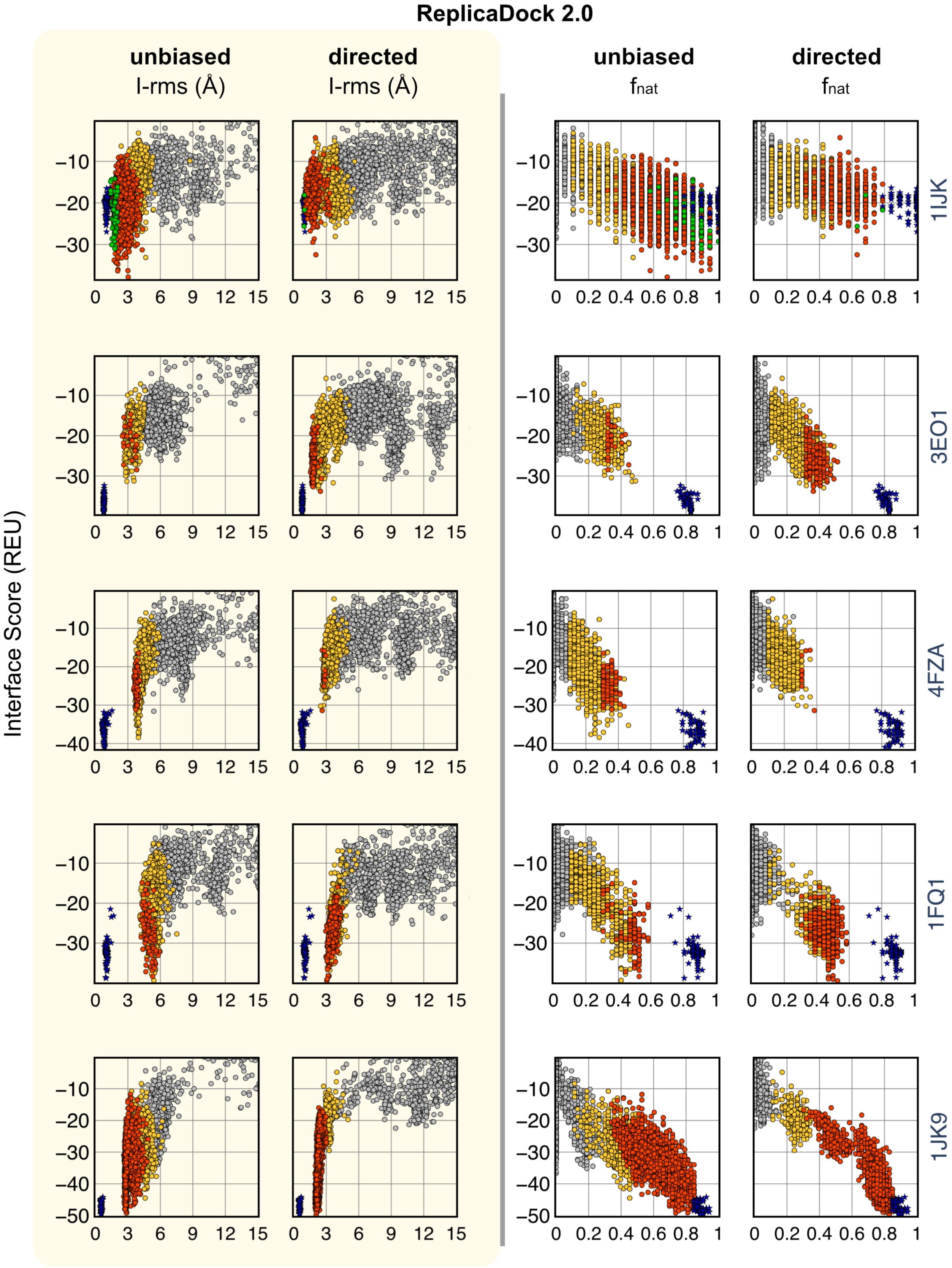
ReplicaDock 2.0
unbiased
I-rms (Å)
directed
I-rms (Å)
unbiased
fnat
directed
fnat
Interface Score (REU)
1IJK
3EO1
4FZA
1FQ1
1JK9

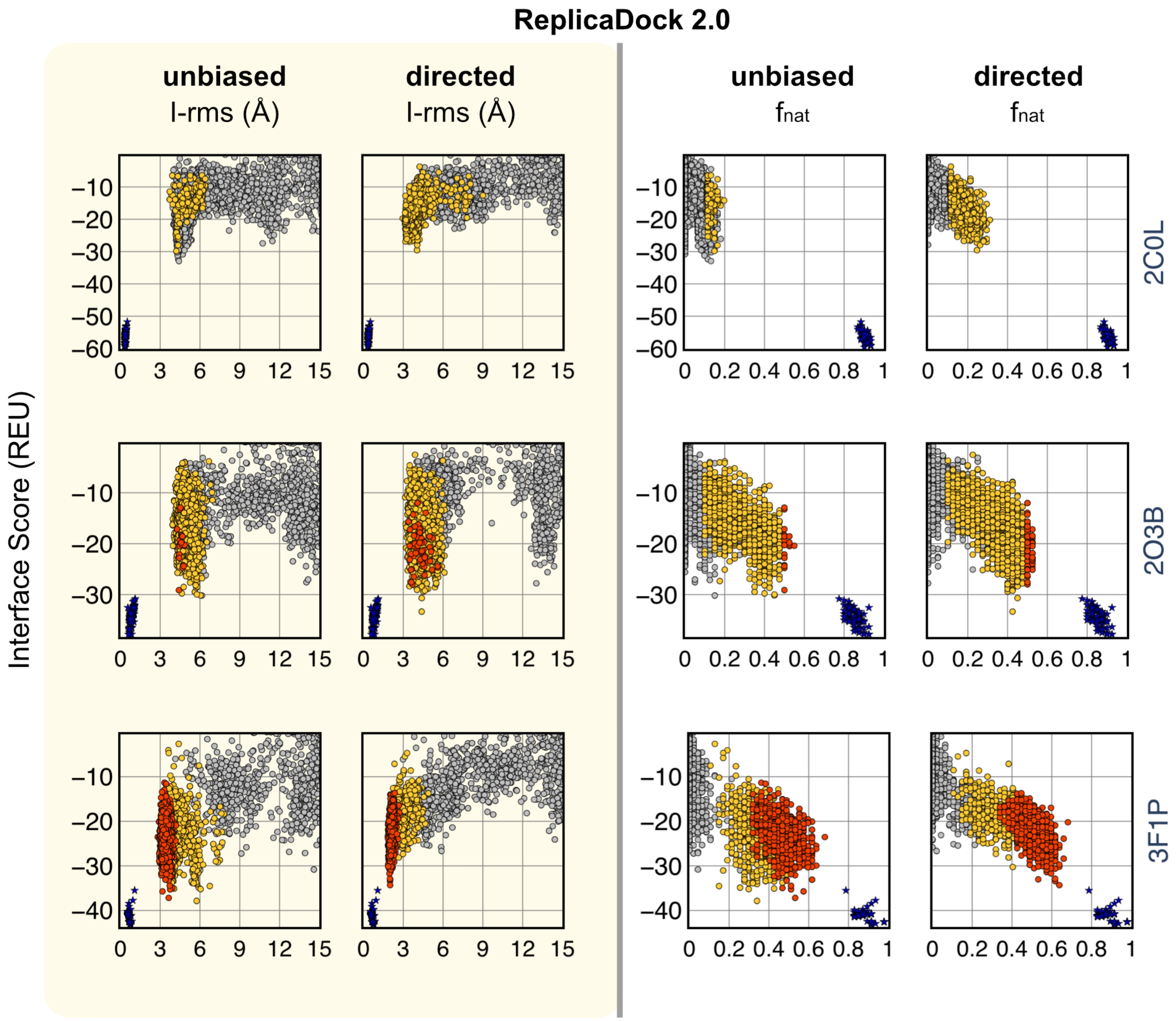


**Fig. S16.** Interface Score versus Interface-RMSD(Å) plots and Interface Score versus $f_{\mathrm{nat}}$ plots after the complete protocol with RosettaDock 4.0 and with directed induced-fit sampling for ReplicaDock 2.0 for benchmark 5.0 targets.
